# Nuclear depletion of RNA binding protein ELAVL3 (HuC) in sporadic and familial amyotrophic lateral sclerosis

**DOI:** 10.1101/2021.06.03.446017

**Authors:** Sandra Diaz-Garcia, Vivian I. Ko, Sonia Vazquez-Sanchez, Ruth Chia, Olubankole Aladesuyi Arogundade, Maria J Rodriguez, Don Cleveland, Bryan J. Traynor, John Ravits

**Affiliations:** Department of Neurosciences, University of California, San Diego, La Jolla, CA 92093-0670; Department of Cellular and Molecular Medicine, University of California, San Diego, La Jolla, CA 92093-0670; Ludwig Institute for Cancer Research, University of California at San Diego, La Jolla, CA, USA; Laboratory of Neurogenetics, National Institute on Aging, National Institutes of Health, Bethesda, MD 20892-3707

**Keywords:** ALS, neurodegeneration, ELAVL3, TDP-43, RNA binding proteins, C9orf72, SOD1, loss of function

## Abstract

Amyotrophic lateral sclerosis is a progressive fatal neurodegenerative disease caused by loss of motor neurons and characterized neuropathologically in almost all cases by nuclear depletion and cytoplasmic aggregation of TDP-43, a nuclear RNA binding protein (RBP). We identified ELAVL3 as one of the most downregulated genes in our transcriptome profiles of laser captured microdissection of motor neurons from sporadic ALS nervous systems and the top dysregulated RBPs. Neuropathological characterizations showed ELAVL3 nuclear depletion in a great percentage of remnant motor neurons, sometimes accompanied by cytoplasmic accumulations. These abnormalities were common in sporadic cases with and without intermediate expansions in ATXN2 and familial cases carrying mutations in C9orf72 and SOD1. Depletion of ELAVL3 occurred at both the RNA and protein levels and a short protein isoform was identified but it is not related to a TDP-43-dependent cryptic exon in intron 3. Strikingly, ELAVL3 abnormalities were more frequent than TDP-43 abnormalities and occurred in motor neurons still with normal nuclear TDP-43 present, but all neurons with abnormal TDP-43 also had abnormal ELAVL3. In a neuron-like cell culture model using SH-SY5Y cells, ELAVL3 mislocalization occurred weeks before TDP-43 abnormalities were seen. We interrogated genetic databases but did not identify association of ELAVL3 genetic structure associated with ALS. Taken together, these findings suggest that ELAVL3 is an important RBP in ALS pathogenesis acquired early and the neuropathological data suggest it is involved by loss of function rather than cytoplasmic toxicity.

## Introduction

Amyotrophic lateral sclerosis (ALS) is a fatal neurodegenerative disease characterized by adult-onset progressive motor impairments from degeneration of upper and lower motor neurons^1–3^. Ninety percent of cases are sporadic (sALS) and their causes are unknown. Ten percent of cases are familial (fALS), and over 65 genes have been identified, the most common genes being C9orf72 and SOD1^4,5^. The signature neuropathological hallmarks of ALS are nuclear depletion, mislocalization and cytoplasmic aggregation of TDP-43 and these are characteristic in 97% of cases--essentially all sALS and most fALS except for SOD1 and FUS mutated ALS^6^. The discovery of TDP-43, a nuclear RNA binding protein (RBP), identified the importance of RNA biology in ALS pathogenesis and led to identification of several other RBPs including FUS^7–9^, TAF15^10,11^, hnRNPA1^12–16^, hnRNPA2B1^17^, EWS^18^, and MATR3^19^. It is unclear how these RBPs contribute to pathogenesis and the debate has focused on loss of their nuclear functions highlighted neuropathologically by nuclear depletion and cytoplasm toxicity highlighted neuropathologically by cytoplasmic accumulation of inclusions and aggregations^5,20–23^.

While ALS is a disease that specifically affects motor neurons in the central nervous system, many of the altered RBPs involved in ALS are ubiquitously expressed. ELAVL (or Hu) proteins are RBPs highly expressed in neurons and fundamental for the central nervous system development. They were discovered as antigens from patients with paraneoplastic neurological syndromes^24,25^. Four ELAVL family members–ELAVL1 (HuA or HuR), ELAVL2 (HuB), ELAVL3 (HuC), and ELAVL4 (HuD)—have been defined. ELAVL1 is ubiquitously expressed in many cell types and ELAVL2, ELAVL3, and ELAVL4 are specifically expressed in peripheral and central neurons throughout development and therefore are sometimes referred to as neuronal ELAVLs (nELAVLs)^26–29^. ELAVL proteins modulate mRNA stability, splicing, and translational efficiency by binding AU-rich elements (ARES) or alternative polyadenylation sites^26,30–33^. The ELAVL proteins are involved with neuronal, axonal and synaptic structure including maturation, differentiation, maintenance, and survival^30,34,35;26,34,36–40^. While the recognition of the role of nELAVLs proteins in general and ELAVL3 in particular in neurological diseases has mainly focused on paraneoplastic neurological syndromes, they have also been previously implicated in ALS pathogenesis^24,25,39,41,42^.

Because of the importance of RBPs in ALS pathogenesis, we analyzed RBPs in our two independent studies profiling transcriptional changes in sALS laser captured motor neurons^43,44^. We identified ELAVL3 because of the degree and statistical significance of the downregulation, the reproducibility of its downregulation in two independent data sets, its exclusively neuronal expression, and its relatively underrecognized role in ALS. We performed extensive neuropathological characterizations and comparisons to TDP-43 pathology. We found several abnormalities in ELAVL3: reduced RNA levels, nuclear depletion, scarce cytoplasmic aggregation, depletion of normal full-length protein, and a short isoform in reduced quantity. The abnormalities were even more prevalent than TDP-43 abnormalities and, in addition, all neurons with abnormal TDP-43 also had abnormal ELAVL3 but not vice versa. Surprisingly, ELAVL3 was also abnormal in SOD1 mutant ALS, where TDP-43 is known to remain normal. ELAVL3 abnomalities were observed upstream of TDP-43 abnormalities in an *in vitro* cell model. Interrogation of genetic structure did not reveal significant susceptibility. Thus, ELAVL3 is an important RBP that is upstream in ALS pathogenesis.

## Results

### Identification of ELAVL3 as a candidate RBP in sALS pathogenesis

We reviewed our previously published transcriptomic data obtained from laser captured spinal motor neurons in sALS nervous systems^43,44^. We focused our attention on candidate RBPs and identified three that were abnormally expressed in both studies—down regulation of ELAVL3 and upregulation of HNRPA1 and NOL8. Of these, ELAVL3 was the most significantly downregulated in magnitude, statistical significance, and reproducibility (Fig. 1)—in sALS, it’s expression was reduced to 42% of normal (log2 fold change −1.2581 with p =0.0009)^43,44^ or 44% of normal (log2 fold change −1.1831 with p=4.00E-08)^43,44^ (Fig. 1A and B). The other RBPs that were differentially expressed in both data sets were slightly upregulated: hnRNPA1 (log2 fold change 1.22, p=0.02^44^ and log2 fold change 1.01396, p=0.016^43,44^ and NOL8 (log2 fold change0.20, p=0.055^44^ and log2 fold change 0.56, p=0.001^43,44^ (Suppl. Table 3). Interestingly, other RBPs including TARDBP, FUS, TAF15, hnRNPA2B1, EWS, and MATR3 were not differentially expressed. We validated the RNA levels of ELAVL3 and TARDBP in frozen spinal cord tissues different from those tissues used in our published reports using qPCR (3 control and 7 sALS) and again found significant downregulation of ELAVL3 (log 2-fold change 0.04, p= 0.01) and not TARDBP (log2 fold change 0.34, p=0.92) (Fig. 1D and E). These supported that our findings in the laser capture microdissection studies are reproducible and consistent.

**Figure 1.**
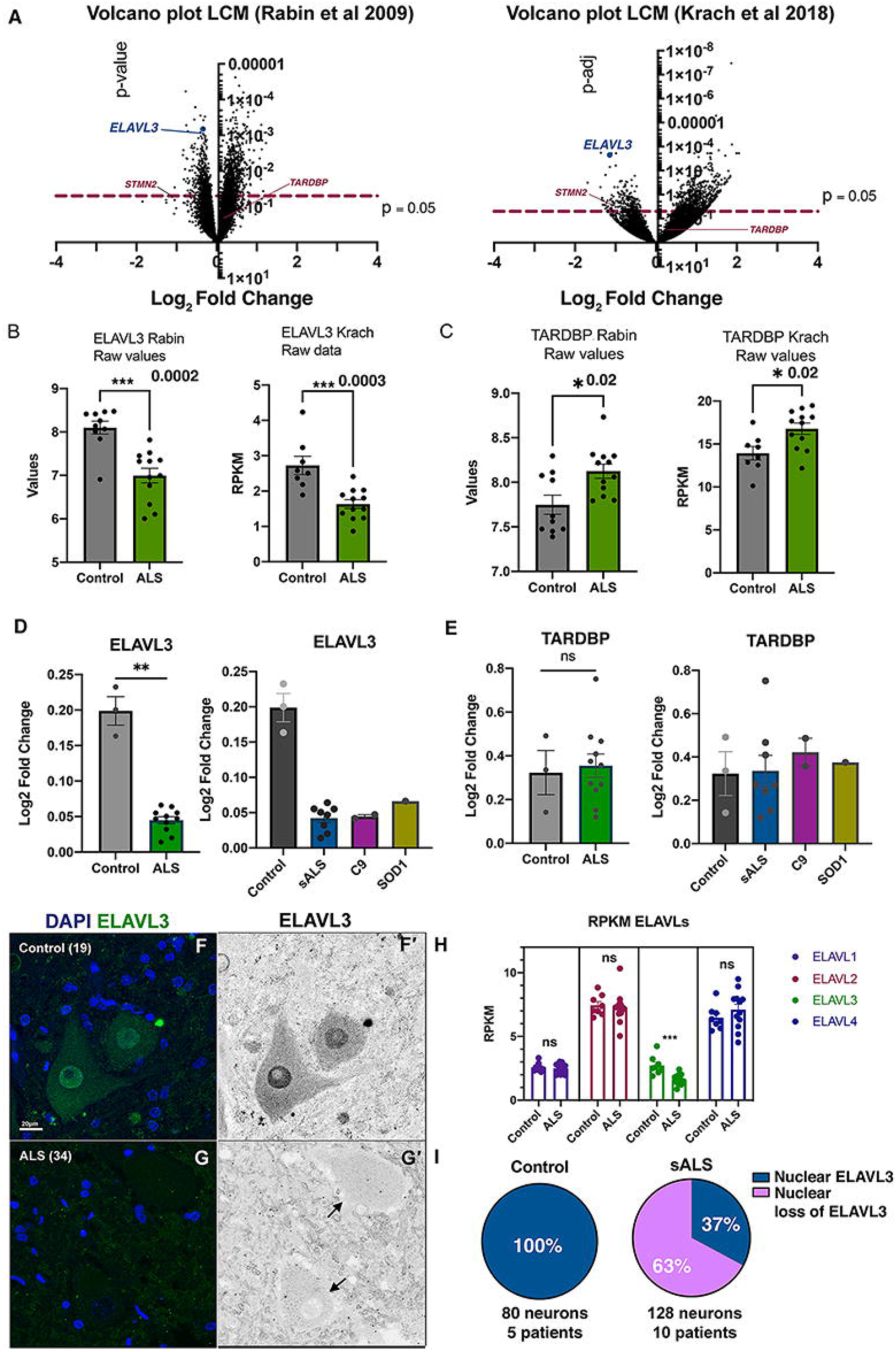
ELAVL3 downregulation in sALS motor neurons. A. Volcano plots of RNAs from laser captured motor neurons comparing sALS and control showing significant *ELAVL3* downregulation in two independent studies. B-C. Raw expression values for *ELAVL3* (B) and *TARDBP* (C). D-E. Validation by qPCR of bulk tissue (left) and by ALS subcategories (right) of *ELAVL3* (D) and *TARDBP* (E). F-G. Immunofluorescence of motor neurons of human spinal cords in control (F) and sALS (G). H. Pie chart of frequency of ELAVL3 nuclear location and depletion. I. Expression of other *ELAVLs* family members. [Scale bar 20μm].

### ELAVL3 mislocalizes from the nucleus

We studied ELAVL3 neuropathologically using IHC and IF (Fig. 1 and Suppl. Fig. 1). As predicted from the transcriptome expression data, in sALS, there were striking abnormalities of ELAVL3 protein expression in alpha motor neurons (Fig. 1G-I and Fig. 2E-H). In control nervous systems, ELAVL3 was clearly observed in nuclei and diffusely in the cytoplasm of essentially all spinal alpha motor neurons (Fig. 1F, Fig. 2A-D and Fig. 3A-C). ELAVL3 was exclusively neuronal, and we did not observe nuclear or cytoplasmic presence in glial cells. In sALS, abnormalities had two main neuropathological phenotypes: depletion of normal nuclear ELAVL3 (Fig. 1G and Fig. 2E-H) and cytoplasmic accumulations, mainly as dot-like inclusions and only rarely as thread-like or fibrillar-like skeins (Fig. 2F and Fig. 3I-H, respectively). ELAVL3 was also observed as diffuse staining along axons and dendrites (Suppl. Fig. 2). While previous studies in a mouse model of ELAVL3 KO showed spheroid formations along the axons in Purkinje cells in cerebellum^26^, we did not observe such formations in axons in anterior horns of sALS patients using MAP2 and TUJ1 markers (Suppl. Fig. 2). ELAVL3 often accumulated in the nucleolus of neurons in which there was ELAVL3 nuclear depletion, interesting in light of our recently reported observation of nucleolar abnormalities in sALS (Suppl. Fig. 3)^45^. We also observed nuclear ELAVL3 depletion from nuclei and rare dot-like aggregates in neurons located in layer V of motor cortex of sALS (Suppl. Fig. 4). Interestingly, in one control nervous system from an individual who carried an ATXN2 intermediate length repeat expansion but did not have neurological disease, we observed rare nuclear depletion of ELAVL3 and dot-like inclusions in a few neurons in the motor cortex but not spinal cord (Suppl. Fig. 5). These findings establish that ELAVL3 is a significant RBP in sALS neuropathology.

**Figure 2.**
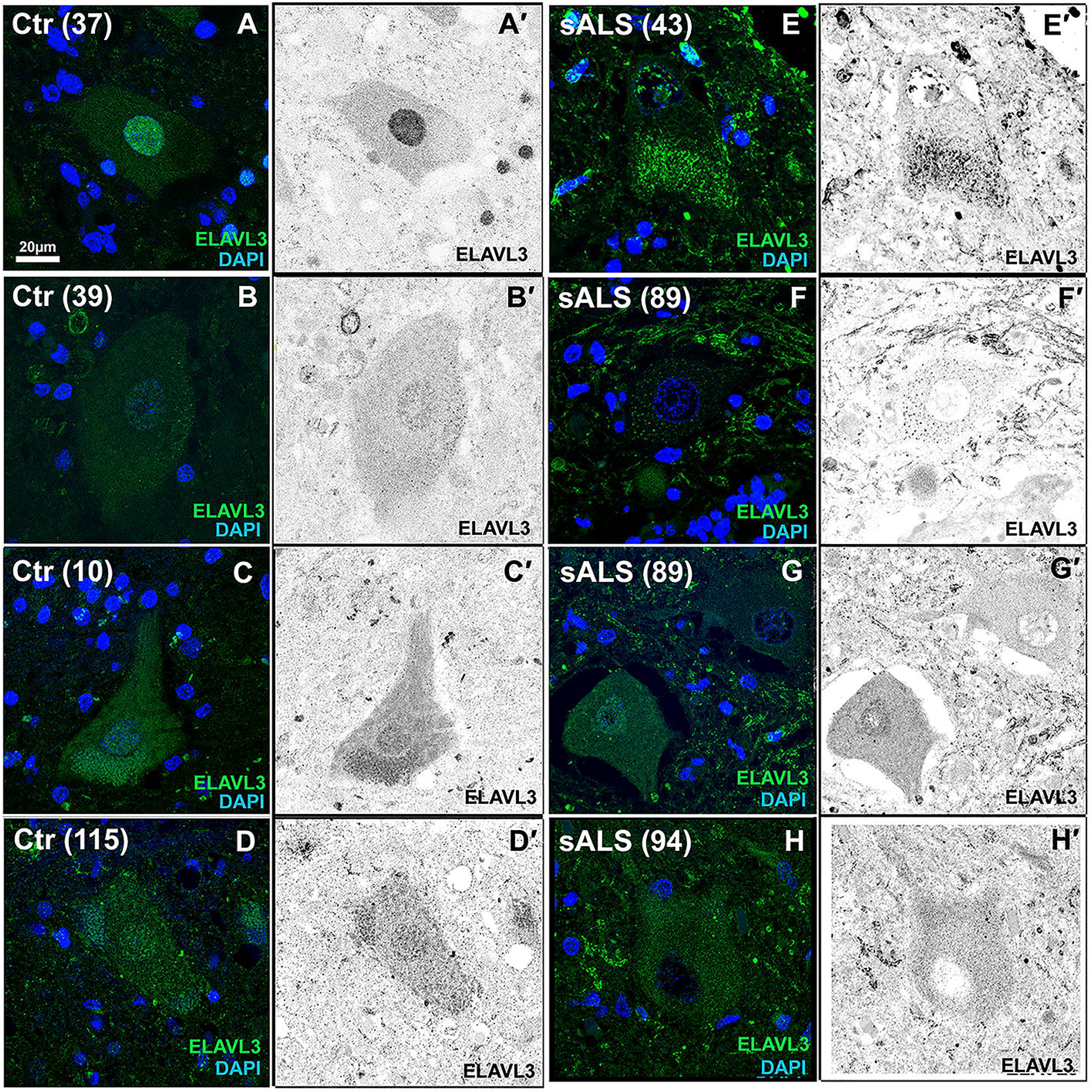
Immunofluorescence of spinal motor neurons in sALS. A-D. Control motor neurons showing ELAVL3 is nuclear and slightly cytoplasmic. E-H. SALS motor neurons showing ELAVL3 depleted in the nucleus, decreased in cytoplasm, and rare cytoplasmic inclusions (F). [ELAVL3 green, DAPI blue, scale bar 20μm].

**Figure 3.**
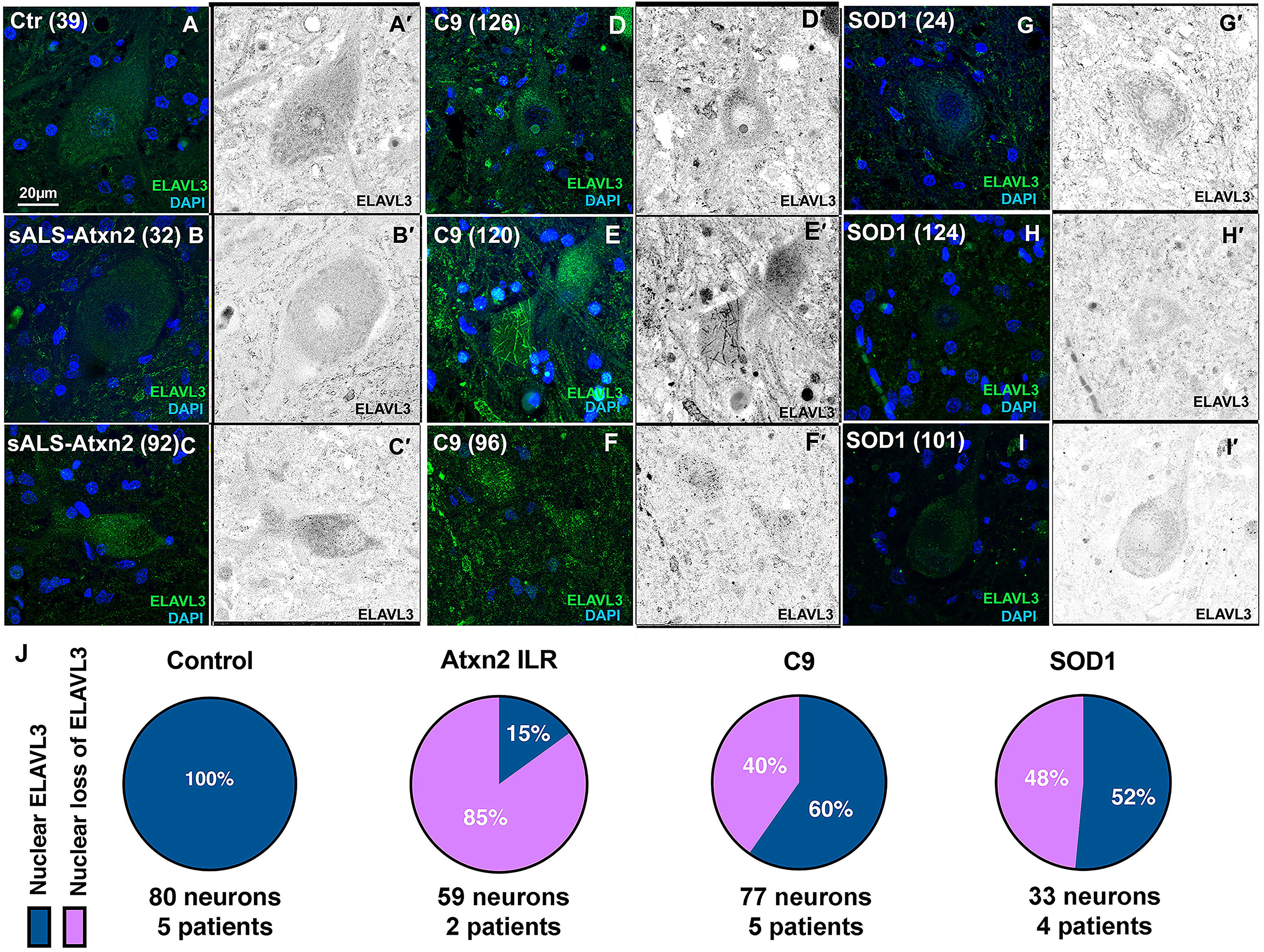
Immunofluorescence of spinal motor neurons in fALS. A. Control motor neurons show nuclear and cytoplasmic ELAVL3. B-C. SALS with intermediate length expansions in ATXN2 showing nuclear depletion and cytoplasmic inclusions. D-F. C9orf72 ALS motor neurons showing nuclear depletion and occasional fibrillary-like (E) and dot-like inclusions (F) are seen. G-I. SOD1 ALS motor neurons showing nuclear depletion and dot-like inclusion (I). J. Pie chart of frequency of ELAVL3 nuclear location and depletion in fALS. [ELAVL3 green, DAPI blue, scale bar 20μm].

### Nuclear depletion of ELAVL3 rather than cytoplasmic aggregation is characteristic

We quantified ELAVL3 abnormalities in 161 motor neurons from 11 sALS nervous systems in the spinal cord using IF. In this, we included only motor neurons that had observable nuclei as identified by DAPI staining (Table 1). The average number of such motor neurons was 15 in each nervous system and the range was 6-35. Overall, 114 (71%) of the motor neurons examined had depletion of nuclear ELAVL3 and 48 (42%) had cytoplasmic accumulations of ELAVL3 (Fig. 1G and 2E-H) (Table 1). Of the 114 motor neurons that had nuclear depletion, 48 (42%) also had cytoplasmic accumulations of ELAVL3 and 66 (58%) did not. Of the 48 motor neurons that had cytoplasmic accumulations of ELAVL3, all 48 had nuclear depletion. These findings suggest that ELAVL3 nuclear depletion is more significant than cytoplasmic accumulations in sALS (p<0.00001 using Chi square analysis and Suppl. Table 4).

**Table 1:** sALS key pathological findings (n=11 nervous systems, 161 neurons)

|  |  |  |
| --- | --- | --- |
|  | ELAVL3 | TDP-43 |
| Neurons with nuclear depletion, total | 114 (71%) | 74 (46%) |
| •Neurons with nuclear depletion that have cytoplasmic inclusions | 48 (42%) | 54 (73%) |
| •Neurons with nuclear depletion that don't have cytoplasmic inclusions | 66 (58%) | 20 (27%) |
| Neurons with cytoplasmic inclusions, total | 48 (30%) | 59 (37%) |
| •Neurons with cytoplasmic inclusions that also have nuclear depletion | 48 (100%) | 54 (92%) |
| •Neurons with cytoplasmic inclusions that don't nuclear depletion | 0 (0%) | 5 (8%) |

### ELAVL3 abnormalities are also seen in fALS nervous systems including SOD1 mutant ALS

We We compared ELAVL3 in nervous systems with known genetic contribution and unknown genetic contribution. 85% of 59 motor neurons examined in two sALS nervous systems with intermediate length repeat expansions of ATXN2 (Fig. 3B-C) and 40% of 77 motor neurons examined in five C9ORF72 fALS had depletion of nuclear ELAVL3 (Fig. 3D-G). While it is well-recognized that nuclear depletion and cytoplasmic aggregation of TDP-43 do not occur in SOD1 ALS^6^, surprisingly we found that 48% of 33 residual motor neurons examined in four SOD1 ALS nervous systems also had nuclear depletion of ELAVL3 (Fig. 3H-J). Additionally, we found ELAVL3 but not TARDBP RNA levels appeared to be down in frozen spinal cord tissues from 2 C9orf72 and 1 SOD1 nervous systems (Fig. 1D). These results support our observation of depleted ELAVL3 in familial cases in tissue samples. Finally, we examined ELAVL3 in three ALS-related mouse models—TDP-43^ΔNLS^ ^46^, C9orf72^47^, and SOD1 (SOD1-G93A)--and observed ELAVL3 nuclear depletion in the motor cortex, aligning with our observations in human samples (Suppl. Fig. 6).

### Reduced ELAVL3 and short isoform is present in sALS and fALS

To confirm the downregulation of ELAVL3 observed transcriptionally and by immunofluorescence, we measured ELAVL3 protein levels by immunoblotting in sALS, C9 and SOD1 ALS cases (Fig. 4). Control samples had a prominent band of ELAVL3 at ~39 kDa corresponding to ELAVL3 full length protein. In every ALS case, we observed a consistent clear depletion of full length ELAVL3. Strikingly, a lower band of ~36 kDa appeared in sALS and fALS cases, the appearance of which was associated with the decrease of 39 KDa band, but at ~1/3 the abundance seen in non-ALS tissue (Fig. 4B). To ensure that ELAVL3 depletion was not due to neuronal loss in ALS tissue, we also measured neuronal marker TUJ1 and confirmed that the decrease of ELAVL3 at protein level was independent of lack of neurons in the tissues. This shorter form of ELAV3 is calculated to be 25-30 amino acids shorter than the full-length form. Knowing that a cryptic exon in intron 3 of ELAVL3 was identified by knocking down TDP-43 in a cellular model^48^, we sequenced from exon 3 to exon 4 in sALS and control spinal cord tissue samples and did not find either a cryptic exon or a difference (data not shown). In addition, we interrogated RNA-seq expression data and did not find differences ^43,49^ (Suppl. Fig. 7). For interest, other differentially expressed genes in ALS whose missregulation could be related to either TDP-43 or ELAVL3 mRNA binding are shown in Supplementary Table 5-8. To determine if this shorter isoform might be related to post-translational cleavage such as at caspase and calpain cleavage, we interrogated known databases and protein cleavage tools and did not identify an obvious site that would result in 25-30 amino acid shortage. Thus, the exact nature of the short isoform is unclear and it might be a product of degradation.

**Figure 4.**
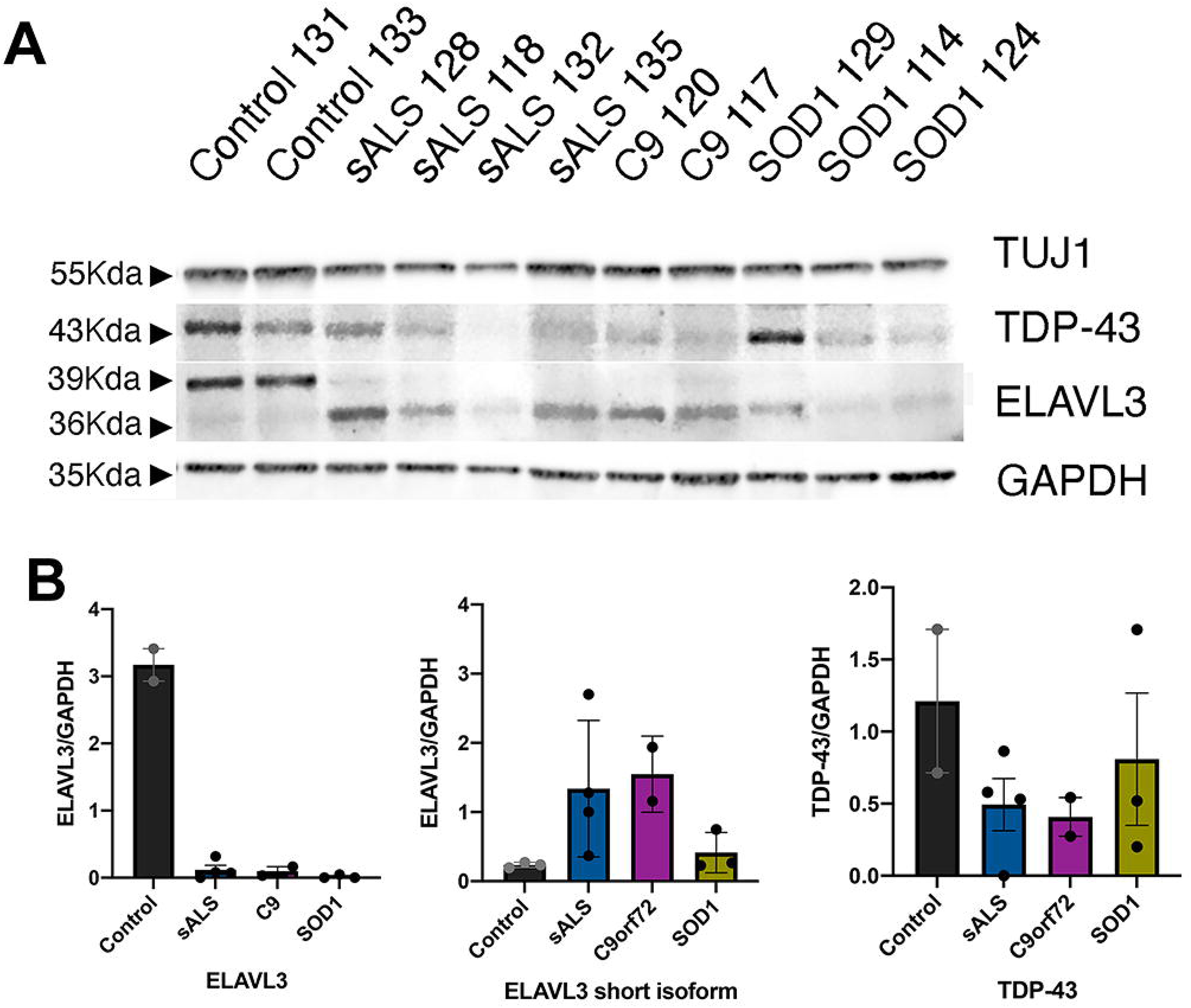
Immunoblotting of spinal cord in sALS and fALS. A. Western blot of healthy donors, sALS and fALS spinal cords for neuronal marker TUJ1, TDP-43, ELAVL3 and GAPDH. B. Quantifications of ELAVL3 and TDP-43 normalized to GAPDH in western blot. Original uncropped blots are shown in Supplementary Figure SX.

### ELAVL3 abnormalities are more prevalent and earlier than TDP-43 abnormalities

We evaluated TDP-43 in the same 161 motor neurons from the 11 sALS nervous systems using co-IF. TDP-43 was well visualized in the nuclei of neurons and glia cells in control tissue, and sALS patients tissue exhibited the expected mix of TDP-43 nuclear depletion and cytoplasmic accumulations, especially aggregates (Table 1 and Suppl. Fig. 8). Overall, 74 (46%) of the motor neurons had depletion of nuclear TDP-43 and 59 (37%) had cytoplasmic accumulations of TDP-43. Of the 74 motor neurons that had depletion of nuclear TDP-43, 54 (73%) also had cytoplasmic accumulations of TDP-43 and 20 (27%) did not. Of the 59 motor neurons that had cytoplasmic accumulations of TDP-43, 54 (92%) also had nuclear depletion. Nuclear depletion and cytoplasmic aggregates were not significantly different (p=0.09 using Chi square analysis and Suppl. Table 4). This supports current consensus that TDP-43 sALS neuropathology has both nuclear depletion and cytoplasmic accumulation and contrasts to ELAVL3, where nuclear depletion is the hallmark neuropathologic change.

We then directly compared ELAVL3 and TDP-43 abnormalities in the same neurons. Neurons with nuclear depletion of ELAVL3 were more frequent than TDP-43 abnormalities and encompassed all TDP-43 pathology (Suppl. Table 4). 46% of neurons with nuclear depletion of ELAVL3 had abnormal TDP-43 (either nuclear depletion or cytoplasmic accumulation), but 100% of neurons with pathological TDP-43 also had nuclear depletion of ELAVL3 (Fig. 5). In addition, we found almost all cytoplasmic pathological inclusions did not colocalize (Suppl. Fig. 9). We compared ELAVL3 and TDP-43 in the fALS nervous systems with ATXN2 intermediate expansions and C9ORF72, and observed depletion of nuclear ELAVL3 even when TDP-43 was nuclear (Fig 5E). Importantly, in nervous systems from SOD1 mutations, we again observed ELAVL3 abnormalities (33% of neurons) and did not observe TDP-43 pathology in ELAVL3-abnormal neurons, where TDP-43 was indeed nuclear (Fig 5E and F).

**Figure 5.**
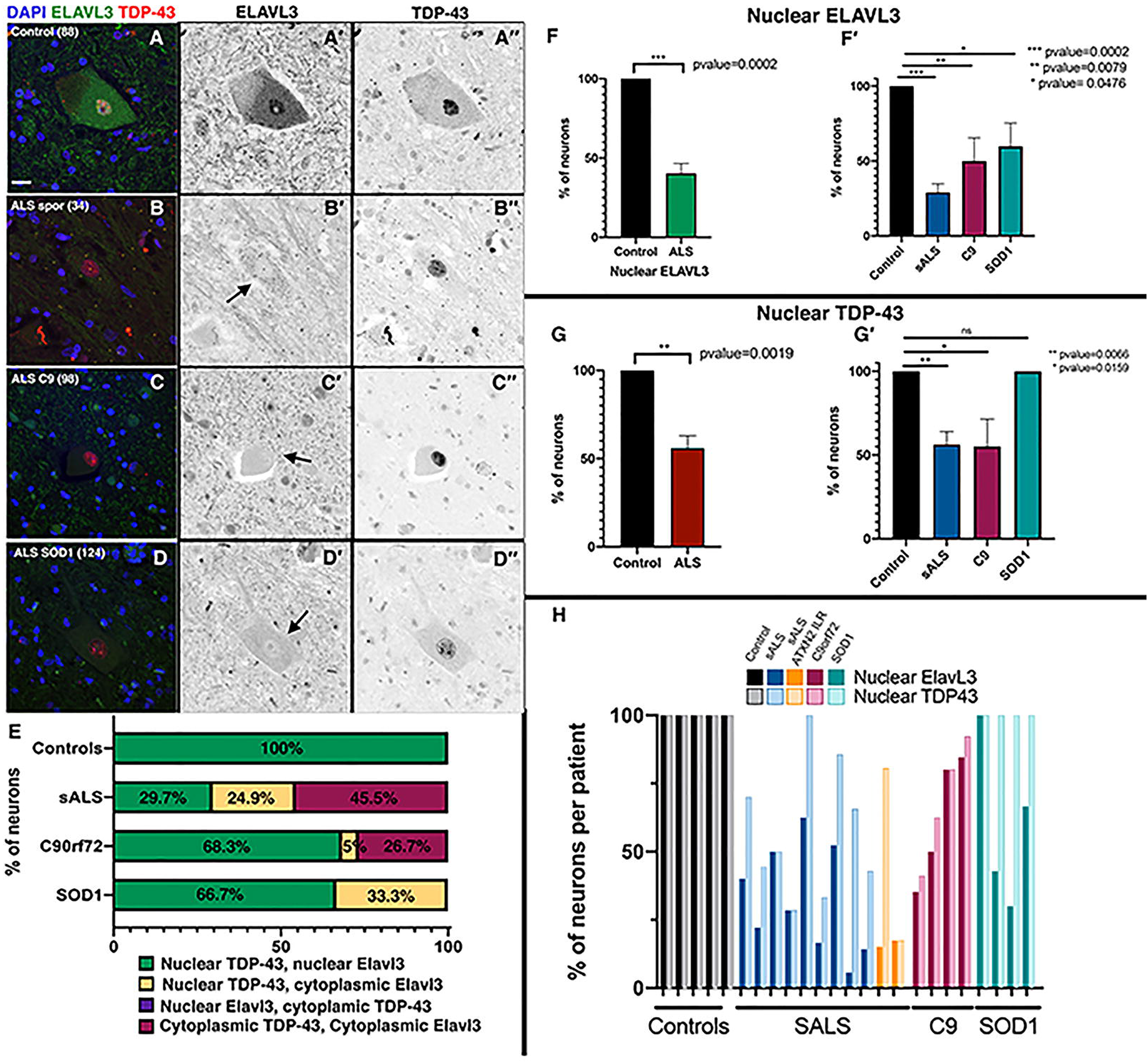
ELAVL3 abnormalities are upstream of TDP-43. Double-labeled immunofluorescence of ELAVL3 and TDP-43 in spinal motor neurons. A. Control motor neuron showing ELAVL3 is present in the nucleus and diffuse in the cytoplasm and TDP-43 is nuclear. B-D. Motor neurons from sALS (B), C9orf72 (C), and SOD1 (D) show ELAVL3 nuclear depletion is observed when TDP-43 is nuclear (B-D). E. Statistical comparison of ELAVL3 and TDP-43 abnormalities in same neurons. F. Statistical analysis of ELAVL3 abnormalities in ALS overall and in sorted groups. G. Statistical analysis of TDP-43 abnormalities in ALS overall and in sorted groups. H. Comparison and variance of ELAVL3 and TDP-43 abnormalities in each nervous system sorted by groups. [ELAVL3 green, TDP-43 red, DAPI blue, scale bar 20μm].

### ELAVL3 nuclear depletion is upstream of TDP-43 in a cellular model

Since our results suggest that ELAVL3 abnormalities may be upstream of TDP-43 abnormalities, we used a neuron-like cellular model under prolonged stress to study the temporal dynamics including nuclear depletion, cytoplasmic localization, and inclusion formation of both proteins. We applied Rapamycin, a common ER stressor^50^, to cell cycle-arrested SH-SY5Y cells which had endogenous TDP-43 tagged with GFP^50^. One day after the stress was applied, ELAVL3 was mislocalized and formed cytoplasmic inclusions while TDP-43 remained predominantly nuclear (Fig. 6). TDP-43 remained normal until 4 weeks, at which time it was depleted in the nucleus and formed dot-like inclusions in the cytoplasm. This model supports that ELAVL3 abnormalities occur earlier and are upstream of TDP-43 abnormalities.

**Figure 6.**
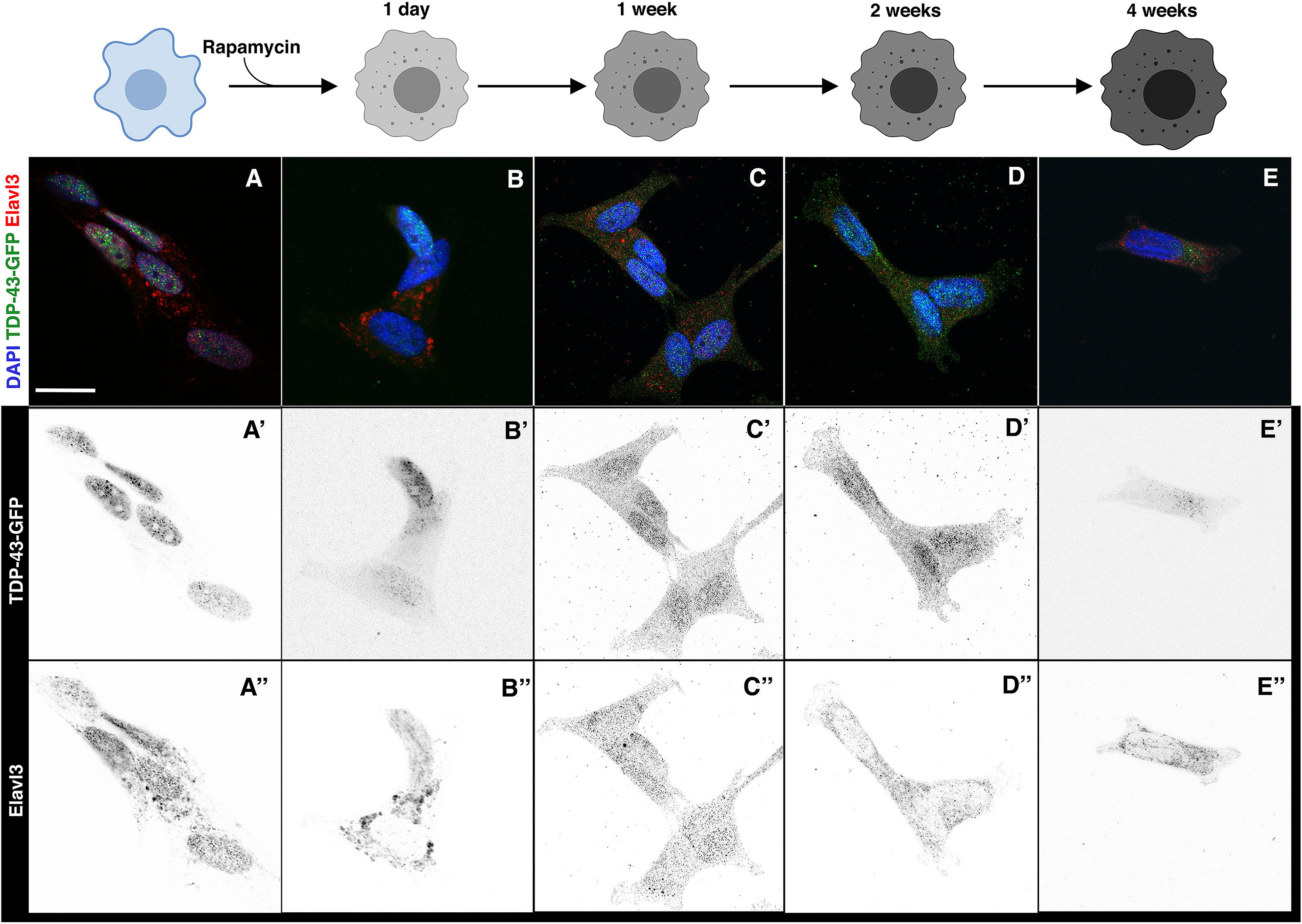
Nuclear clearance of ELAVL3 is prior to TDP-43 mis-localization in cellular model. Cell-cycle arressted SH-SY5Y cells are stressed by rapamycin and TDP-43 and ELAVL3 are studied over time at unstressed (A), 1 day (B), 1 week (C), 2 weeks (D) and 4 weeks (E) after after rapamycin treatment. ELAVL3 abnormalities appear early and TDP-43 late.

### Other ELAVL family members show variable and infrequent pathology

Because ELAVLs family members have high sequence homology and may have redundant functions in the nervous system, we analyzed the expression pattern of ELAVL1 (Ctr=2 and sALS=6), ELAVL2 (Ctr=4 and sALS=10) and ELAVL4 (Ctr=6 and sALS=31). ELAVL1 was expressed in the nuclei and diffusely in the cytoplasm of neurons and glial cells in control nervous systems (Fig. 7). In sALS, C9 and SOD1 nervous systems, nuclear expression of ELAVL1 was usually similar to controls but more variable expressed, and rare cytoplasmic inclusions were seen, occasionally colocalizing with TDP-43 speckles or encapsulating TDP-43 aggregates in the cytoplasm (Fig. 7 and Suppl. Fig. 10). ELAVL2 was normally expressed in the nuclei and diffusely in the cytoplasm of neurons but not glial cells in control nervous systems. In sALS and C9 nervous systems, ELAVL2 was variably reduced in the nucleus and colocalized with fibrillar aggregates of TDP-43 in the cytoplasm (Fig. 7). We did not observe ELAVL2 abnormalities in SOD1 nervous systems. ELAVL4 was variably expressed in the nucleus and cytoplasm of neurons but not glia in controls. In sALS, C9 and SOD1 nervous systems, nuclear expression of ELAVL4 was more variable expressed than controls, and cytoplasmic inclusions were seen usually as dot-like inclusions occasionally colocalizing with TDP-43 (Fig. 7 and Suppl. Fig. 10). To compare all 4 proteins to each other and establish variability in the same nervous systems, three nervous systems were examined for all four ELAVLs (Ctr=1, sALS=1 and SOD1=1). We found only ELAVL3 was consistently and significantly abnormal (Table 2 and Suppl. Fig 10). Taken together, these findings suggest that ELAVL3 is unique within the ELAVL family, but other ELAVL family members have cytoplasmic alterations in the presence of TDP-43 abnormalities.

**Figure 7.**
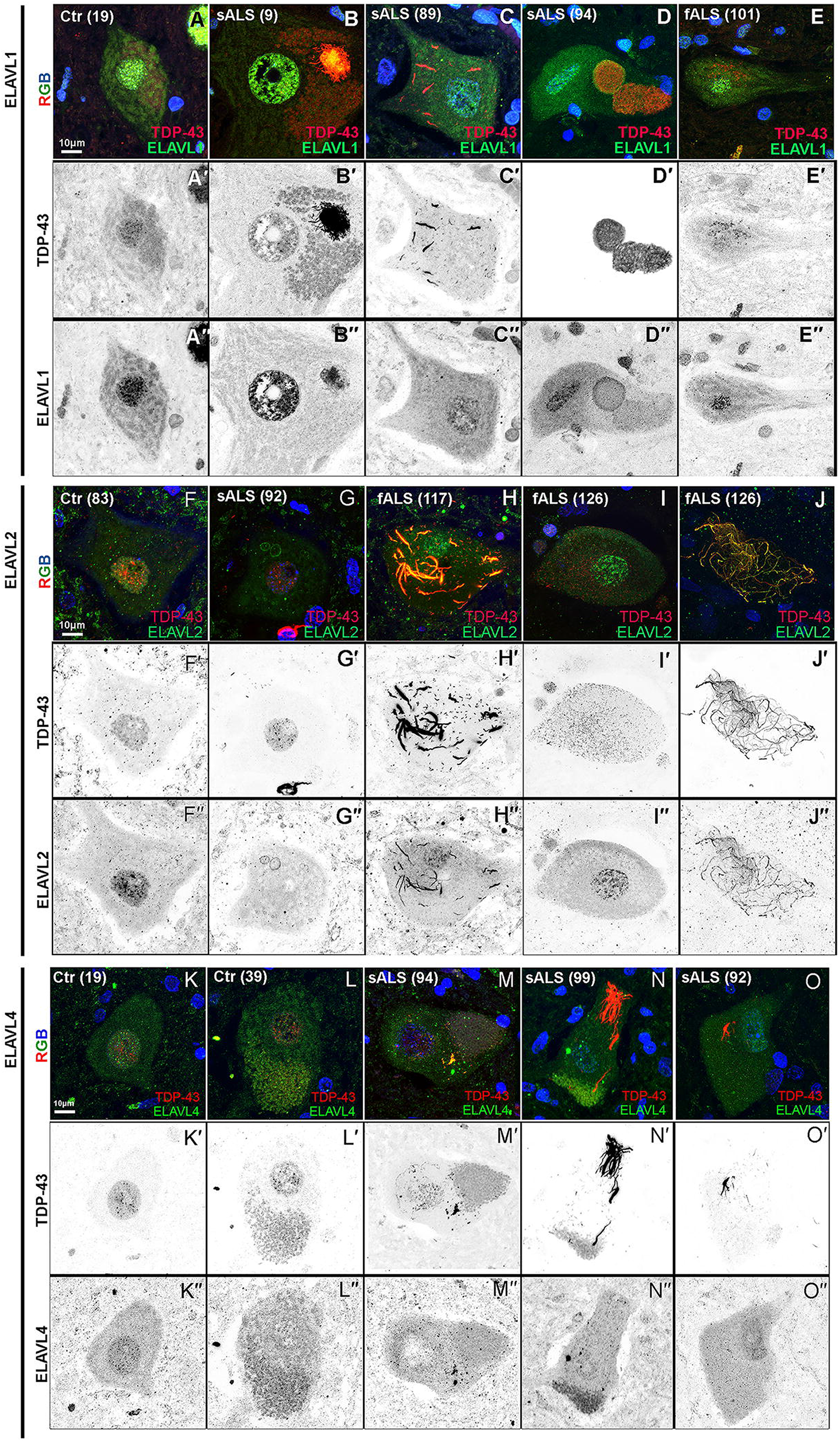
ELAVL family members are also frequently abnormal but only in presence of TDP-43 aggregates. A-E. ELAVL1 in control (A), sporadic, (B-D) and SOD1 (E). Nuclear ELAVL1 expression is maintained or slightly reduced and, in the cytoplasm, and can be seen either colocalizing or encapsulating TDP-43 aggregates or forming its own inclusions. F-J. ELAVL2 expression in control (F), sporadic (G), and C9orf72 (H-J). Nuclear ELAVL2 expression is maintained or reduce depending on the ALS sample and in the cytoplasm can be seen either colocalizing TDP-43 fibrillary aggregates (H-J), but not with speckles (I). K-O. ELAVL4 in control (K-M) and sporadic samples (M-O). Nuclear ELAVL4 expression is more diffuse in controls than other ELAVLS and in ALS it is reduced or depleted from the nucleus it variably colocalizes with TDP-43 aggregates in the cytoplasm.

**Table 2:** Summary of ELAVL Family

| Table 2: Summary of ELAVL Family |  |  |  |  |
| --- | --- | --- | --- | --- |
| In presence of TDP-43 pathology | ELAVL1 | ELAVL2 | ELAVL3 | ELAVL4 |
| NUCLEUS | No change/<br>Decrease | No change/<br>Decrease | **Nuclear loss** | No change/<br>Decrease |
| CYTOPLASM | No change/<br>Decrease | No change/<br>Decrease | No change/<br>Decrease | No change/<br>Decrease |
|  | Rare dot-like inclusions | No dot inclusions | Dot like inclusions | Dot like inclusions |
|  | Colocalization with TDP-43 or encapsulation of aggregates | Often colocalize with TDP-43 fibrils but not dot like inclusions | No colocalization with TDP43. Rarely colocalization with large TDP-43 aggregates | Sometimes colocalize with TDP-43 aggregates |

### There are no ELAVL genetic variants associated with ALS

To investigate the contribution of common ELAVL variants to disease risk, we performed association testing on 750 common variants present in ELAVL1,2,3 and 4 genes in 10,599 cases and 42,279 controls^51^. There were no variants within these genes that passed the genome-wide threshold of 5 × 10^−8^, implying that these common variants do not have a significant effect on risk at the population level in sALS cases (Suppl. Fig. 11). At the rare variant level, we performed a gene-based analysis on variants that were predicted to be missense or have a loss-of-function effect using the SKAT-O method and did not find excess presence of rare variants in cases compared to controls. From these results, it does not appear to be that there are rare variants in these genes that drive/modify risk for ALS, at least this is true for this set of imputed variants in the tested cohort.

## Discussion

Because of the focal clinical onset of ALS and anatomical propagation of disease over time, the timing of degeneration desynchronizes in motor neurons and they display different degrees of molecular changes neuropathologically^52^. This allowed us to identify upstream candidates using a genomics approach^43,44^ and from this, we identified ELAVL3 as a candidate RBP of interest. Indeed, we found striking neuropathological abnormalities of ELAVL3 in motor neurons of sporadic and familial ALS tissue samples. The abnormalities were of two varieties—nuclear depletion and cytoplasmic accumulation— and of these, nuclear depletion was hallmark. The importance of nuclear depletion is further supported by the fact that ELAVL3 had reduced expression at both the RNA and the protein levels. Thus, the evidence supports that nuclear depletion and loss of function rather than subcellular location or cytoplasmic accumulation and toxicity are key to the mechanism by which ELAVL3 may be involved in pathogenesis. The neuropathological differences between cytoplasmic accumulation and nuclear depletion that we observed are unlikely due to sampling, since the cytoplasmic accumulations are diffusely scattered in the cytoplasm and the cross-sectional areas of cytoplasm are significantly greater than cross-sectional areas of nucleus, which would bias in the opposite direction. The difference is also unlikely to be due to the detection threshold of antibody since it is equal in either subcellular location.

It is not unreasonable to think ELAVL3 pathobiology in ALS may be different from TDP-43. In TDP-43 pathobiology, the preponderance of evidence indicates subcellular mislocalization rather than expression is key. Both nuclear depletion and cytoplasmic accumulation of TDP-43 usually appear together neuropathologically and our neuropathological data indicate both nuclear loss and cytoplasmic accumulation are significant. TDP-43 expression is tightly regulated and much data support that it is not differentially expressed although our own transcriptome data if anything suggest slight upregulation. Unlike TDP-43, ELAVL3 does not have a canonical nuclear localization signal (NLS) or a low complexity glycine rich prion domain and it is significantly down regulated at the RNA and protein levels. Our data further suggest that ELAVL3 abnormalities are upstream of TDP-43 in the process of motor neuron degeneration. This is supported by three lines of reasoning. First, all neurons with TDP-43 abnormalities displayed ELAVL3 abnormalities but many neurons with ELAVL3 abnormalities displayed normal nuclear TDP-43. Thus, TDP-43 pathology was a subset of ELAVL3 pathology, not vice versa. Second, ELAVL3 abnormalities were also seen in SOD1 nervous systems, where TDP-43 abnormalities are not present. The role SOD1 plays in sALS is debated. Regardless, it is accurate to say that ELAVL3 pathology is widespread and prevalent in sALS and fALS from the two most common mutations SOD1 and C9orf72. Neither SOD1 or TDP-43 are as clearly generalizable neuropathologically as ELAVL3 is, although it is possible that TDP-43 or SOD1 are abnormal at a molecular level that are not visible neuropathologically. Third, our cell culture data in which cell cycle arrested non-dividing neuronlike TDP-43-GFP SH-SY5Y cells were artificially stressed clearly showed ELAVL3 abnormalities (mislocalization and inclusion formation) long before TDP-43 abnormalities (mislocalization and inclusion formation), suggesting a more sensitive and prompter response to stress. These results were recently shown in detail in an experimental study reporting uniform downregulation of ELAVL3 in extensive ALS iPSC-MNs models at both RNA and protein levels and, importantly, this decrease was present beginning in early development, suggesting ELAVL3 is an early modulator of downstream disease^39^. Taken together, these findings are consistent with an upstream effect of ELAVL3 on both SOD1 and TDP-43, an effect that could be explained, for example, by regulation of ARES in the 3′UTR, which both proteins are known to have^53–55^.

The short ELAVL3 isoform was an unexpected finding. Since the changes on immunoblotting mobility are of3kD, the abnormal isoform is expected to be 25-30 amino acids shorter than the normal protein. An isoform of ELAVL3 is known to exist, but it is only differs by 6 amino acids different (isoform 2 is missing amino acids 251-257). Since we used an antibody to exon 2 epitopes near the N-terminal in both the neuropathological and biochemical studies, it seems reasonable to conclude that the shortening is after exon 2. One possibility is inclusion of a cryptic exon from reduction of TDP-43 such as has been identified in STMN2^48^ and UNC13A^58,59^. Indeed, a cryptic exon due to TDP-43 downregulation has been identified at intron 3 of ELAVL3^48^. However, we could not identify such in the ALS samples sequencing from exon 3 to exon 4 as well as mapping our RNA seq data^43,48^. This is consistent with our other findings that ELAVL3 abnormalities are independent of TDP-43 mechanisms. The possibility that ELAVL3 protein might undergo cleavage was not supported by caspase or calpain cleavage sites predictions in protein cleavage databases. Thus, at this time we are unable to further explain the structure and mechanism of the short isoform.

Previous literature has already implicated ELAVL3 in ALS pathogenesis, although it has received little direct attention and is underrecognized. In a systematic survey of 213 proteins harboring RNA recognition motifs in a yeast functional screen, ELAVL3 (but not the other ELAVLs) had a high toxicity score and, as stated, did not harbor a prion domain, unlike TDP-43, FUS and other RBPs^42^. In human neuronal cell model, TDP-43 and FUS depletion both showed changes in ELAVL3, but they were oppositely regulated: down-regulated by siTDP-43 and up-regulated by siFUS^60^. In a study of TDP-43 loss of function in human motor neurons differentiated from human pluripotent stem cells, ELAVL3 was significantly downregulated and, as mentioned above, a cryptic exon was identified at intron 3^48^. Also as mentioned above, a recent study of iPSC-derived motor neuron models of familial and sporadic ALS found down-regulation of ELAVL3 expression was an early ALS hallmark that began during development and persisted into end stage^39^. Other ELAVLs family members have also been identified in ALS biology and are important in light of neuropathological characterizations of ELAVL2 and ELAVL4, which showed cytoplasmic accumulation and colocalization with TDP-43 aggregates in motor neurons of ALS spinal cords. ELAVLs were shown to be involved as positive gene regulators of SOD1 by binding to ARES in the 3′UTR^55^. ELAVL2 was differentially expressed in motor axons and NMJ, although the findings showed upregulation^35^. In iPSC-derived MN models of FUS mutant ALS, mutant FUS bound ELAVL4 3′UTR, resulting in increased production of the ELAVL4 protein, which were found in cytoplasmic aggregates in the spinal cord of ALS^61^. ELAVL4 has also been shown to be involved in spinal muscular atrophy, another motor neuron disorder^62^.

The suggestion that ELAVL3 abnormalities are upstream in the process of neurodegeneration led us to question if genetic factors might be related to its mis-regulation. However, no significant genetic association of ELAVL3 or other ELAVL family members could be identified, indicating mis-regulation of ELAVL3 is likely an early acquired abnormality. In summary, we have compelling evidence implicating ELAVL3 upstream in sporadic, C9orf72 and SOD1 ALS pathogenesis that does not appear to be genetically based. The triggers and upstream regulators and downstream mechanisms in the neurodegeneration cascade remain unknown. Further studies will elucidate mechanisms of this important RBP.

## Materials and methods

### CNS tissues

Human tissues were obtained from the UCSD ALS tissue repository that was created following HIPAA-compliant informed consent procedures approved by Institutional Review Boards (either Benaroya Research Institute, Seattle, WA IRB# 10058 or University of California San Diego, San Diego, CA IRB# 120056). Spinal cord and brain tissues were acquired using a short-postmortem interval acquisition protocol usually under 6 hours. Tissues were immediately dissected in the autopsy suite, placed in labelled cassettes and fixed in neutral buffered formalin for at least 2 weeks before being dissected and paraffin embedded for indefinite storage. For this study, we evaluated from 6 control cases, 31 sALS cases, 5 C9orf72 cases and 4 SOD1 (Suppl. Table 1).

### CNS region and neuronal identification

Motor neurons were identified by their large size, multipolar cytoplasm, presence of lipofuscin, and large nucleus with a less dense DAPI staining in IF studies. In spinal cord, we examined motor neurons in Rexed lamina IX of anterior horn in cervical, thoracic, and lumbar sections. In the cortex, we examined Betz cells in layer V of the cortex. For quantifications, we chose paraffin blocks that we knew had abundant neurons as revealed by standard H&E light microscopy and previous studies revealing abundant TDP-43 pathology.

### Immunohistochemistry (IHC) and immunofluorescence (IF)

Tissue sections were cut from blocks of formalin-fixed paraffin embedded ALS tissue, obtained from the UCSD ALS bank collection. 6μm-thick tissue sections were de-paraffinized through histology grade CitriSolv (two times for 15 min each) and a graded alcohol series (100, 90, 70 and 50% ethanol (vol/vol) for 5 min each). Then we dipped the slides in de-ionized water, permeabilized for 20 min in 1 × PBS with 0.2% Triton X-100, performed antigen retrieval with 1% Tris-based (Vector # H-3301) in a pressure cooker at 120 °C for 20 min. Sections were further blocked with 2% Fetal Bovine Serum (vol/vol, Atlanta Biologicals S11150) and incubated with the primary antibody. Primary antibodies were ELAVL3 (#55047-1-AP from Proteintech and LS□C167722 from LsBio) TDP-43 mouse (#DR1075-100UG, 1:500; calbiochem), TDP-43 Rabbit (#10782-2-AP Rabbit Proteintech), HuR-Antibody (#11910-1-AP, Proteintech), ELAVL4 (#24992-1-AP Proteintech), and ELAVL2 (#67097-1-AP Proteintech). The antibodies for ELAVL1 and ELAVL3 were previously validated by KO studies^63–66^. ELAVL2 and ELAVL4 antibodies were validated by protein BLAST alignment demonstrating unique protein sequences. The ELAVL antigen sequences against which the antibodies were made are in Supplementary Table 2. These were incubated overnight at 4 °C. For IHC, we quenched with 0.6% H2O2 in Methanol for 15 minutes before the permeabilization step and continue with the protocol. For both IF and IHC protocols, after incubation with primary antibody, sections were washed four times in 1X PBS for 5 minutes and blocked with 2% Normal Donkey Serum (Millipore S30-100ml) before incubation with secondary antibodies in 1X PBS, 2% Normal Donkey Serum (vol/vol) for 1 hr at room temperature. For detection of primary antibodies using IF, we used donkey anti-rabbit, anti-mouse Alexa-488, Cy3 conjugated antibodies (Jackson ImmunoResearch) and goat anti-chicken Alexa 633 conjugated antibody (Invitrogen A-21103) at a 1:500 dilution. The slides were then incubated with DAPI (1 μg/ml) followed by a PBS wash. To reduce autofluorescence noise, quenching with 0.1% Sudan Black in 70% EtOH for 15 s was applied prior to coverslip mounting with ProLong Gold Antifade Mountant with DAPI media (Invitrogen). For IHC after blocked with 2% Normal Donkey Serum, we incubated with ImmPRESS® HRP Horse Anti-Rabbit IgG Polymer Detection Kit, Peroxidase (MP-7401 Vector) and developed the antibody with Vector® NovaRED® Substrate Kit, Peroxidase (HRP) (SK-4800 Vector) for 1-2 minutes until the color was noticed.

### Imaging, digital processing, and semiquantitative analysis

Sections from blocks of the spinal cord and motor cortex were stained with the antibodies described above. We analyzed 2 sections of 6μm per patient. A semiquantitative assessment of RBP nuclear depletion and mislocalization or decreased expression in ALS tissues was performed taking digital photographs using the Nanozoomer at 40X and visualizing under 60X objective in either Nikon confocal A1/LANA system and Leica SP8 confocal systems at UCSD’s Microscopy Core facility. Depending on the degree of neurodegeneration of each tissue, we counted as many motor neurons as could be identified as described above. Once neurons were identified, motor neurons were scored based on the presence or absence of nuclear ELAVL3 and TDP-43, being either present or depleted. For illustrations, images were taken at a resolution of 1,024 × 1,024 pixels with a ×60 objective and between 15-20 z stack images (0.25-0.50 um per stack) were collected per frame, and the maximum intensity projection was used to construct the final image series using Fiji software. The .tif files were loaded into Photoshop, where the RGB channels were split. Once the figure was built, each individual channel was inverted to facilitate visualization. The following criteria were used in scoring cytoplasmic accumulations: *inclusions* were small dot like, round, diffusely scattered speckles that are relatively uniformly distributed; *aggregates* were discrete, dense, long, fibrillary or striae-like (or threadlike) accumulations; and cytoplasmic *inclusion bodies* were large, round, dense deposits in the cytoplasm present in low copy number, usually only one per cell.

### Cell culture

Neuroblastoma SH-SY5Y TDP-43-GFP line was obtained from Cleveland Lab. All cells were maintained in high-glucose Dulbecco’s modified Eagle’s medium (DMEM) (Gibco, U.S.A.), supplemented with 10% FBS (Gibco, U.S.A.), 2□mM□L-glutamine (Lonza, cat. no. BE12-719□F),100 units/ml penicillin, and 100 μg/ml streptomycin, in a humidified atmosphere of 5% carbon dioxide at 37°C. SH-SY5Y TDP-43EGFP cells were plated on 8-well chamber (Ibidi) at 25,000 cells per well. After 24 h, cells were arrested in G1 with Palbociclib (Apexbio) and maintained during the whole length of the experiment. Cell culture media was changed every 3 days. In all the experiments, the SH-SY5Y control cells and the experimental were plated and maintained in the same culture time and media conditions with Palbociclib, without and with rapamycin respectively. 10nM of Rapamycin was diluted in 100ul of DMSO to use as stressor. rapamycin was used at a final concentration of 5μM in the media to stress the cells. rapamycin was maintained in the media for 4 weeks. Cells were fixed with 4% PFA in PBS for 10 min at room temperature After two washes with PBS, cells were permeabilized and blocked with blocking solution (0.1% Triton, 2% BSA in PBS) for 1 h at room temperature. Cells were then incubated overnight with the primary antibody in PBS/0.3% Tween 20. The primary antibodies used for staining were: anti-ELAVL3 (ab56574, Abcam; 1:1000) at day1 and after 1, 2 and 4 weeks after rapamycin application. After three washes with PBS, the cells were subsequently incubated with fluorescently labeled secondary antibodies diluted at 1:500 in 1XPBS for 1 h at room temperature. After three washes with 1XPBS, a secondary antibody (anti-rabbit-555 from Jackson) was applied for 1h. We washed the cells 3 times for 5 minutes each with 1X PBS and mounted with ProLong Gold Antifade Mountant with DAPI media (Invitrogen) to preserve the cells for imaging. SH-SY5Y cells imaging was performed on a Ti1 Nikon microscope with a C2 confocal camera at 60-100x magnification.

### Immunoblotting

To characterize ELAV3 expression in spinal cord samples, frozen tissue from donors and patients (Suppl. Table 1), were weighted and homogenized in ice-cold N-PER Neuronal Protein Extraction Reagent (Thermofisher # 87792) with protease and phosphatase inhibitors in 1mL per 100mg ratio. The samples were spun down and the supernatants were subjected to Pierce™ BCA Protein Assay (Thermofisher #23225). Samples were boiled with Laemmli sample buffer for 10 minutes at 90 °C and to normalize the total amount of proteins among samples 35 μg were loaded in SDS-PAGE (12% 1 mm acrylamide gel with 2,2,2-Trichloroethanol) and, transferred into Nitrocellulose membranes (Bio-rad # 1704159) with Trans-Blot Turbo Transfer System (Bio-Rad# 1704150) (7min, 25V, 2.5 A). Membranes were blocked using iBind™ Flex Fluorescent Detection (Invitrogen # SLF2019) and were incubated with primary (1:500 ELAVL3 (#55047-1-AP Proteintech),1:5000 GAPDH (#2794431, Millipore), 1:500 TDP-43 (12892-1-AP Proteintech), 1:1000 TUJ1 (#801202 Biolegend) and secondary antibodies (1:2000 IRDye 800CW Goat anti-Rabbit IgG # 926-32211 1:2000 IRDye 680RD Goat anti-Mouse IgG # 926-68070, LI-COR) using iBind™ Flex Western System (Invitrogen # SLF2000). NewBlot™ Nitro Stripping # 928-40030 LI-COR was use as a stripping buffer between primary antibody incubation when needed. The images were acquired with an Odyssey® Fc Imaging System from Li-COR. ImageStudioLite method was used to measure the signal intensity.

### Animals

All experimental procedures were approved by the Institutional Animal Care and Use Committee of the University of California San Diego. SOD1G93A, C9orf72 mutants mice tissue samples were courtesy of Don Cleveland laboratory. TDP-43^ΔNLS^ {B6;C3-Tg(NEFH-tTA)8Vle Tg(tetO-TARDBP*)4Vle/J} were obtained from The Jackson Lab (Stock# **028412**).

### Gene specific common variant association and gene-based rare variant analyses

Individual genotyped data consisted of 10,599 cases and 42,279 controls from Nicolas et al ^51^. Control sample data came from the dbGaP webportal (accession numbers: phs000001, phs000007, phs000101, phs000187, phs000196, phs000248, phs000292, phs000304, phs000315, phs000368, phs000394, phs000397, phs000404, phs000421, phs000428, phs000454, phs000615, phs000675, phs000801, and phs000869). Samples were processed using a standard quality control pipeline ^51^. Sample exclusion criteria were: (1) excessive heterozygosity rate (exceeding +/− 0·15 F-statistic), a measure of DNA contamination; (2) low call rate (≤ 95%); (3) discordance between reported sex and genotypic sex; (4) duplicate samples (determined by pi-hat statistics > 0·8), and (5) non-European ancestry based on principal components analysis compared to the HapMap 3 Genome Reference Panel. Related samples (defined as having a pi-hat > 0·125) were included for imputation, but one pair member was removed before the association testing.

Variant exclusion criteria were: (1) monomorphic or palindromic SNPs; (2) variants that showed non-random missingness between cases and controls (p-value ≤ 1·0×10^−4^); (3) variants with haplotype-based non-random missingness (p-value ≤ 1·0×10^−4^); (4) variants with an overall missingness rate of ≥ 5·0%, non-autosomal variants (X, Y, and mitochondrial chromosomes); and (6) variants that significantly departed from Hardy-Weinberg equilibrium in the control cohort (p-value ≤ 1·0×10^−10^).

Imputation was performed against the Haplotype Reference Consortium (HRC 2016) reference panel (hg19) via minimac4 using data phased by Eagle v2.4 provided by Michigan Imputation Server (https://imputationserver.sph.umich.edu/). Only variants in ELAVL1,2,3 and 4 genes with imputation quality score (R2) > 0.3 and minor allele frequency > 0.01 were included for association analysis. Covariates that were adjusted for to account for non-disease related cohort stratification were SEX, age at disease onset, PC1, PC2, PC3, PC4, PC6, PC7, PC8. Regional plots for each gene were generated using LocusZoom (http://locuszoom.org/).

Sequence kernel association test - optimized (SKAT-O) analysis was performed on combined missense and loss-of-function mutations present in ELAVL1,2,3 and 4 genes. The analysis collapses all rare variants and tests for difference in the aggregated burden of rare coding variants between cases and controls. Variants were annotated using ANNOVAR (v.2018-04/16) prior to association testing using RVTESTS (v.2.1.0). Variants were included in the analysis if the minor allele frequency was < 1% and minor allele count was >= 3. The Bonferroni threshold for genome-wide significance for missense category was 5.11 × 10^−6^ (0.05 / 9,790 autosomal genes tested), and for missense and loss-of-function category was 3.60 × 10^−6^ (0.05 / 13,907 autosomal genes tested).

### Statistical analyses

Data were analyzed using GraphPad Prism version 7.0a and are expressed as mean ± standard error of the mean. Significance was assessed using the Student *t* test non-parametric Mann Whitney test, and *p* values less than 0.05 were considered significant. Chi square tests used an online calculator (https://www.socscistatistics.com/tests/).

## Abbreviations

ALS: amyotrophic lateral sclerosis
ARES: AU-rich elements
CNS: central nervous system
fALS: familial amyotrophic lateral sclerosis
MN: motor neuron
nELAVLs: neuronal ELAVLs
NLS: nuclear localization signal
RBP: RNA binding protein
sALS: sporadic amyotrophic lateral sclerosis
3′UTR: 3′-untranslated region

## Author Contributions

SDG analyzed the data of previous papers and designed the experiments. MJR did tissue embedding and histology. SDG and MJR performed the staining. VIK did qPCR analysis, imaging and quantifications. VIK provided mouse tissue and performed staining in mice. OAA helped with cellular models and motor-cortex imaging. SVS did immunoblotting. RC and BT evaluated genetic variants. SDG and JR wrote the paper. SDG, VIK, SVS, OAA, MJR, DWC, BJT, RC and JR discussed the results and contributed to the final manuscript.

---

**Supplementary Figure 1.**
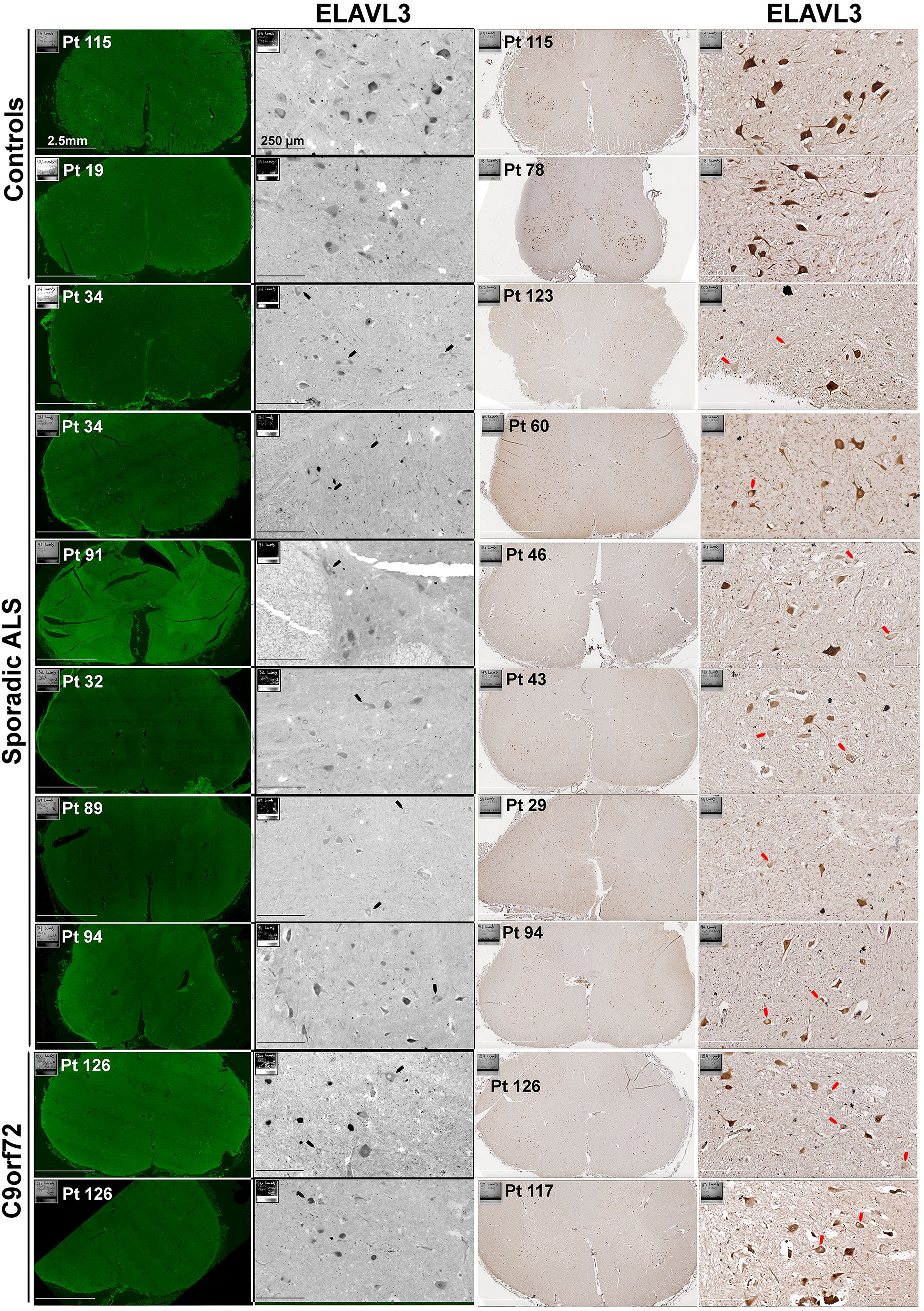
ELAVL3 depletion using two different antibodies and imaging techniques Proteintech antibody is used for IF and LsBio is used for IHC.

**Supplementary Figure 2.**
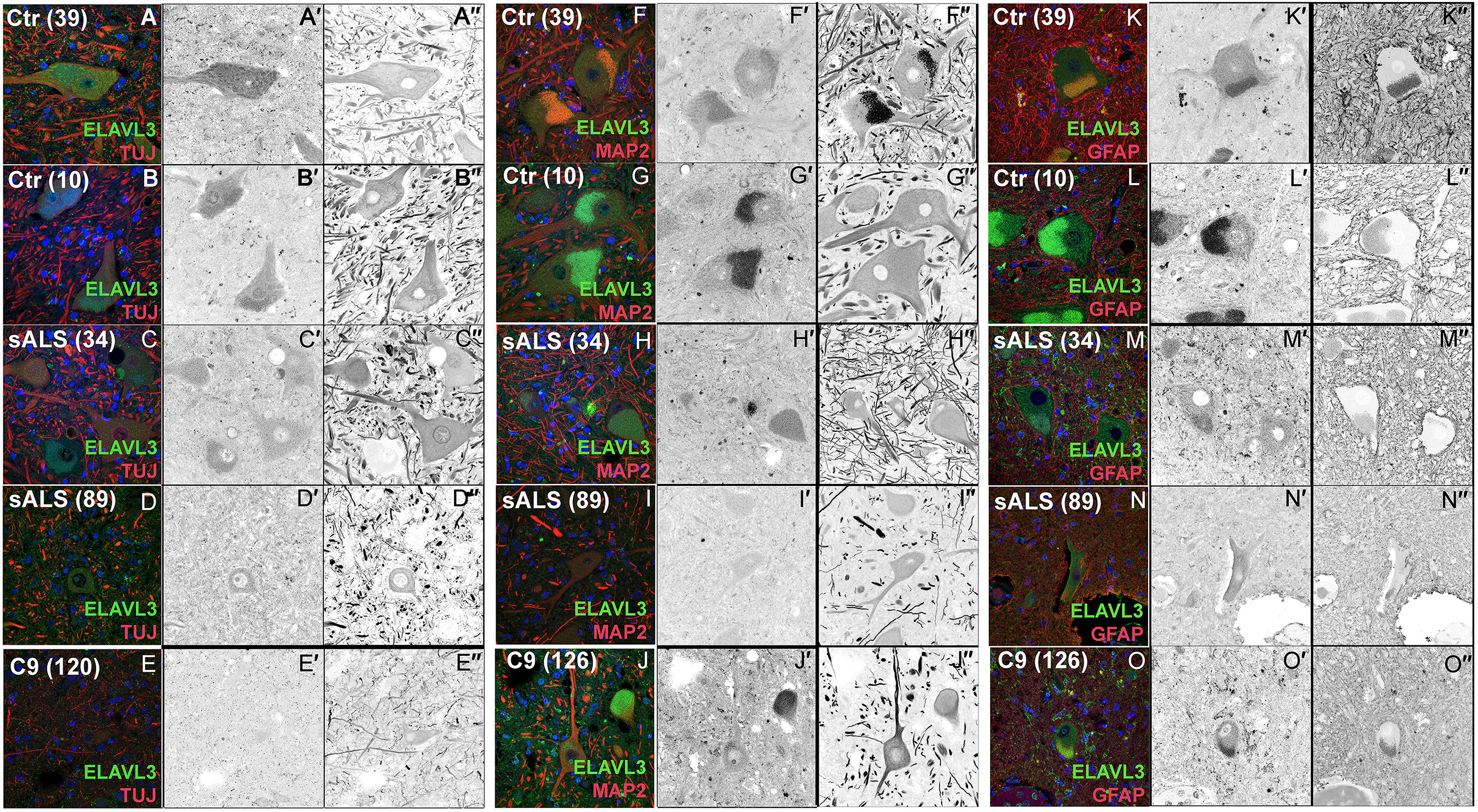
ELAVL3 is expressed in the nucleus and cytoplasm of neurons and can also be seen in axons and dendrites. A-E. Co-staining of ELAVL3 and Tuj1 show no clear accumulation of ELAVL3 in axons in ALS. F-J. Co-staining of ELAVL3 and MAP2 show no clear accumulation of ELAVL3 in dendrites in ALS and rarely in C9orf72. K-O. Co-staining of ELAVL3 and GFAP show no co-localization in ALS.

**Supplementary Figure 3.**
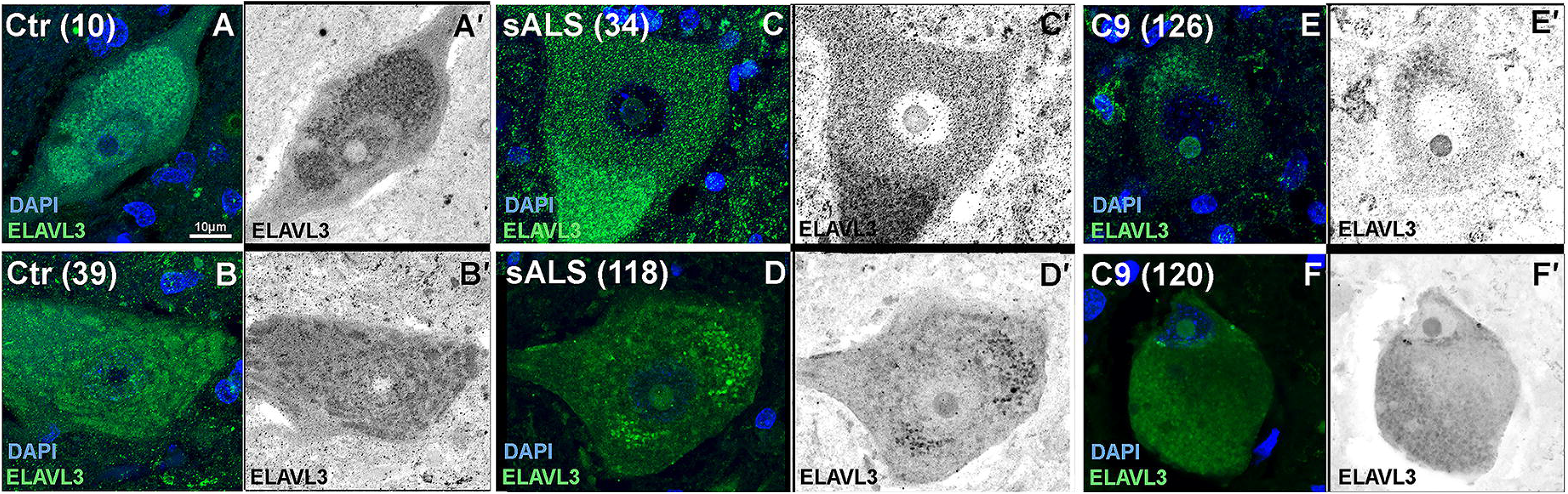
ELAVL3 frequently accumulates in nucleolus of ALS but only rarely in control motor neurons. A-B. Control motor neurons with nuclear ELAVL3 and visible nucleolus without ELAVL3. C-F. ALS motor neurons show nuclear depletion of ELAVL3 and an abnormal accumulation in nucleolus.

**Supplementary Figure 4.**
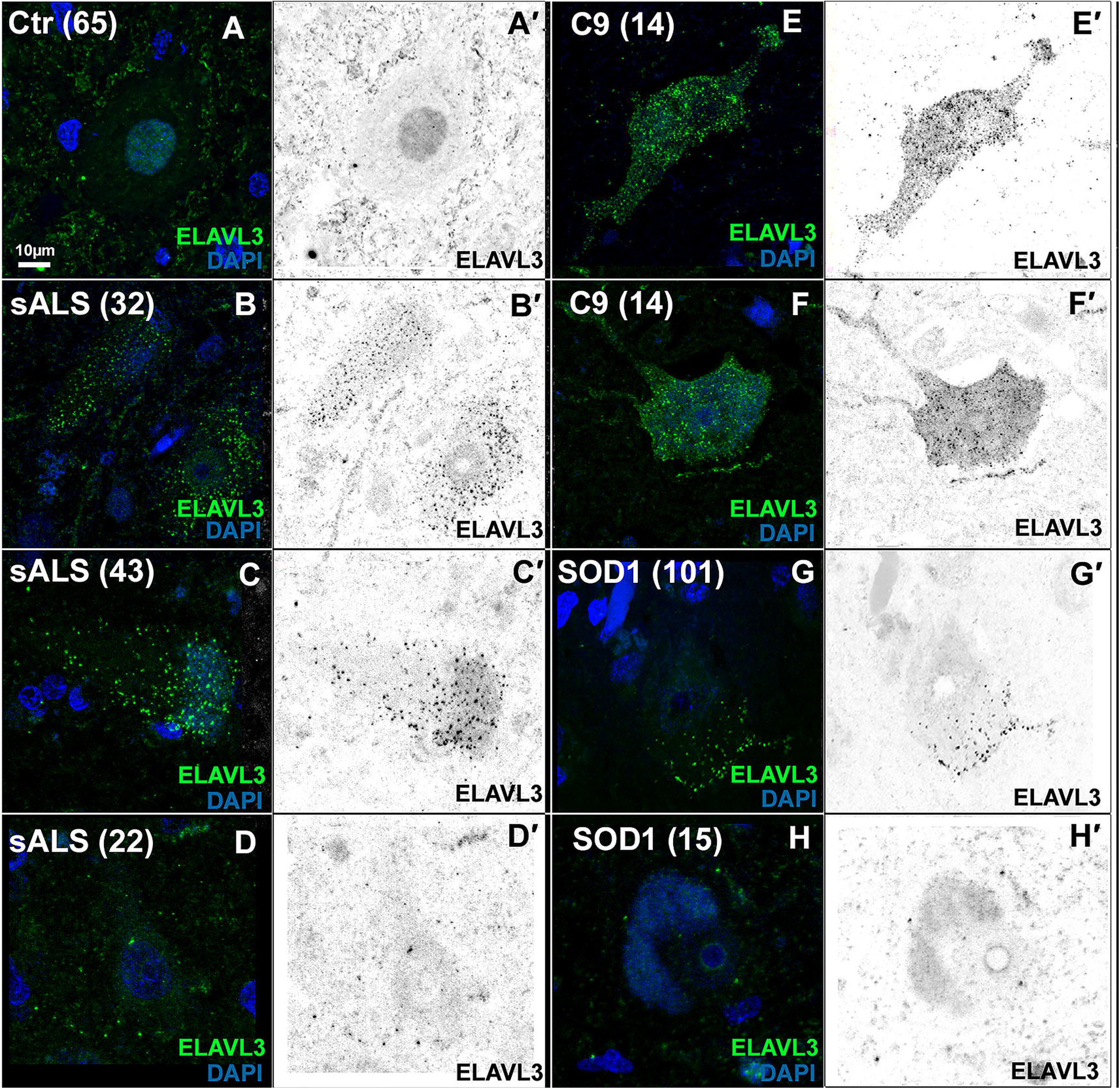
ELAVL3 is mislocalized and forms aggregates in motor cortex of ALS patients. A. Control motor neurons where ELAVL3 is nuclear. (B-D): Sporadic patients motor neurons can have nuclear ELAVL3 (lower neuron in panel B) but also nuclear depleted ELAVL3 as well as cytoplasmic inclusions (B-D). (E-F): C9orf72 motor neurons from motor cortex also have nuclear ELAVL3 (F), nuclear clearance (E) and cytoplasmic inclusions (E-F). (G-H): SOD1 motor neurons from motor cortex also can have nuclear depletion of ELAVL3 (G-H), and cytoplasmic inclusions (G).

**Supplementary Figure 5.**
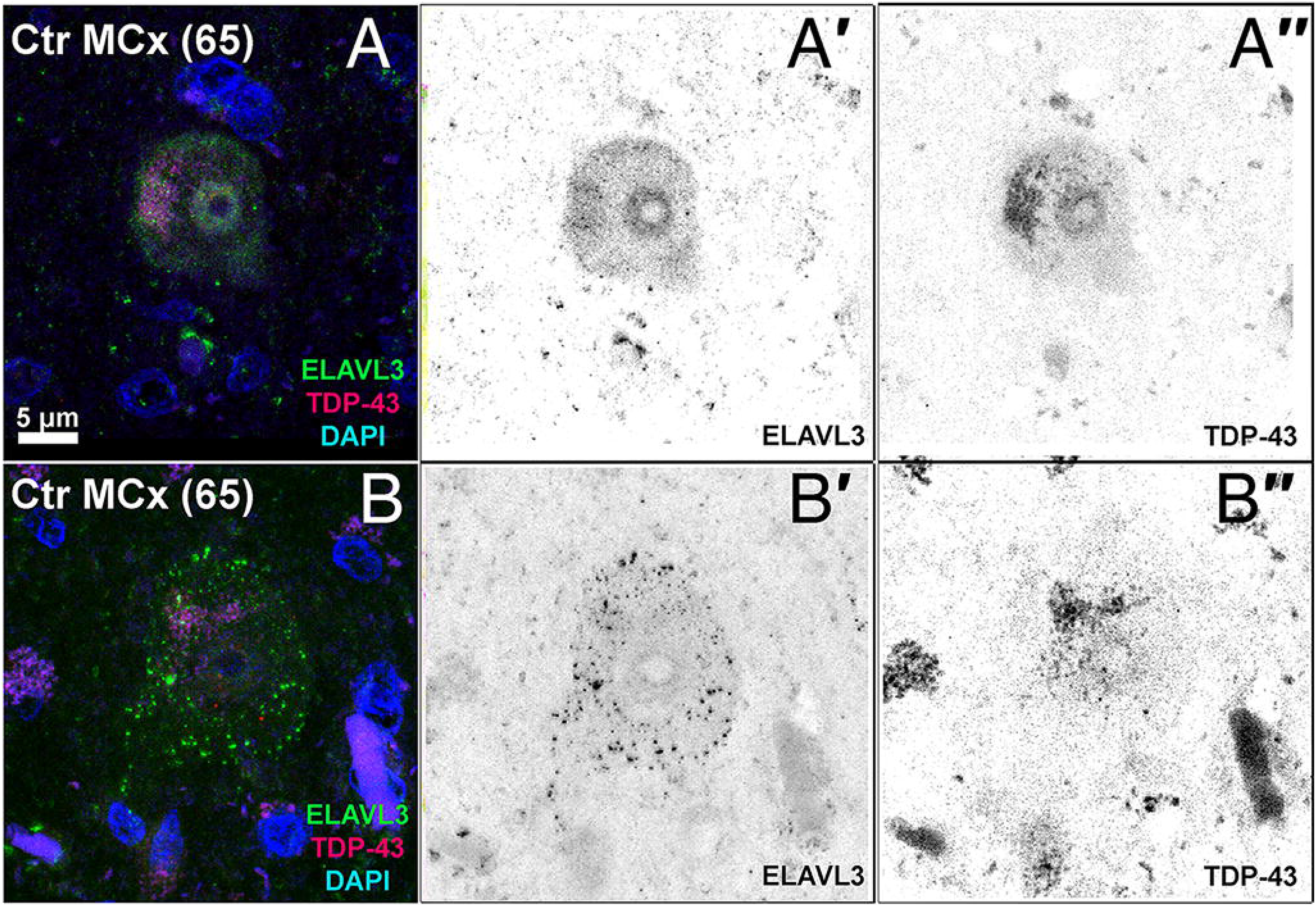
Interesting control patient with rare ELAVL3 abnormalities. (A-B): A few neurons at motor cortex from control 65 that carried intermediate length repeat expansions of ATXN2 had ELAVL3 inclusions in the cytoplasm.

**Supplementary Figure 6.**
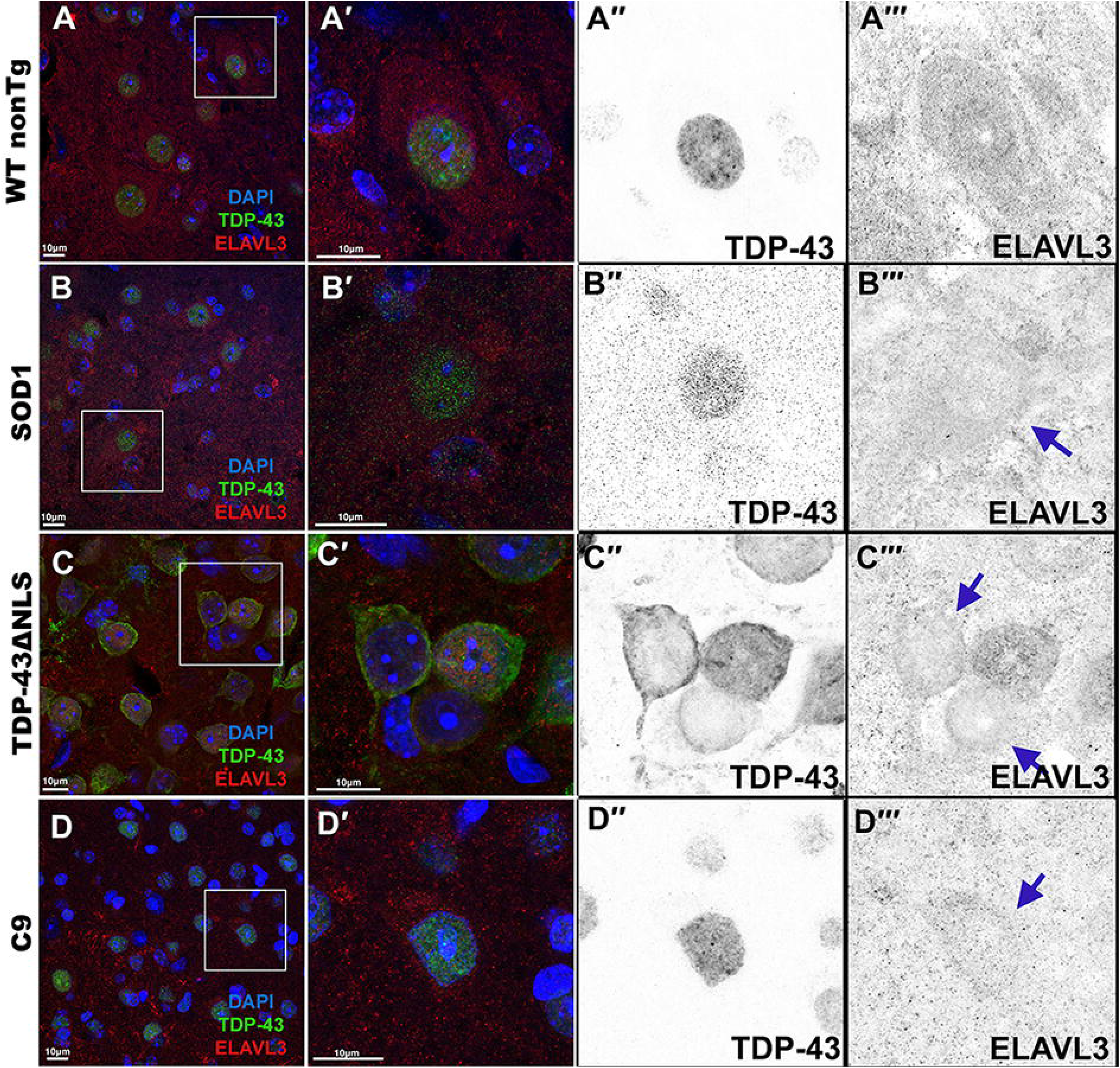
ALS mouse models show nuclear depletion of ELAVL3 in motor neurons. (A): Non-transgenic mice show nuclear ELAVL3 and TDP-43 in motor neurons. (B): SOD1 transgenic mutant mice have lack of nuclear ELAVL3 but normal nuclear TDP-43. (C): TDP-43^ΔNLS^ transgenic mutant mice have lack of both nuclear ELAVL3 and TDP-43 and in addition have cytoplasmic TDP-43 aggregates. (D): C9orf72 mutant mice have lack of nuclear ELAVL3 but have normal nuclear TDP-43. [Scale bar 10μm]

**Supplementary Figure 7.**
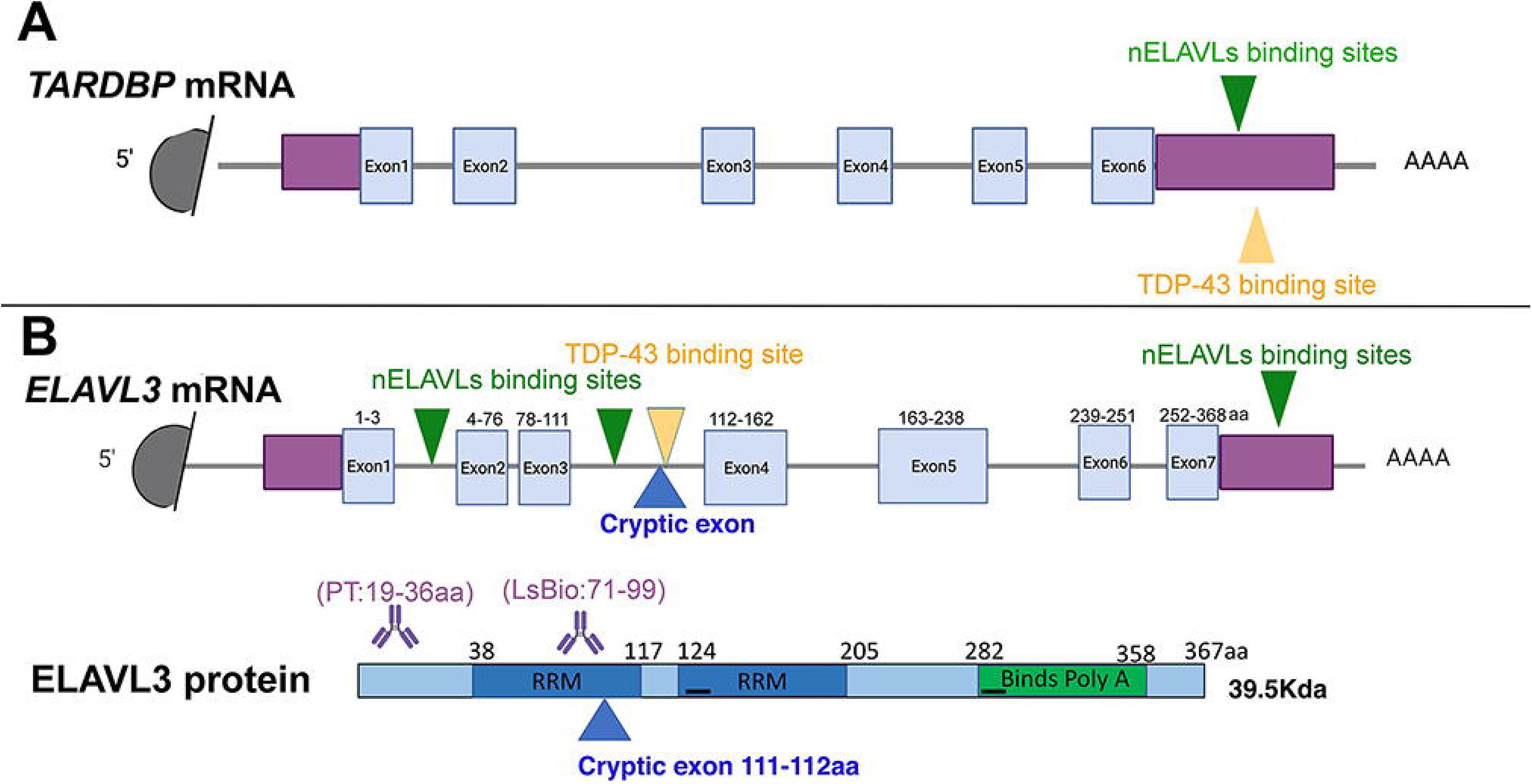
No TDP-43-dependent cryptic exon in ELAVL3. (A) mRNA binding sites in *TARDBP* by nELAVLs and TDP-43, showing binding is at 3′UTR. (B) mRNA binding sites in *ELAVL3* by nELAVLs and TDP-43, showing binding in introns and location of TDP-43-dependent cryptic exon in intron 3.

**Supplementary Figure 8.**
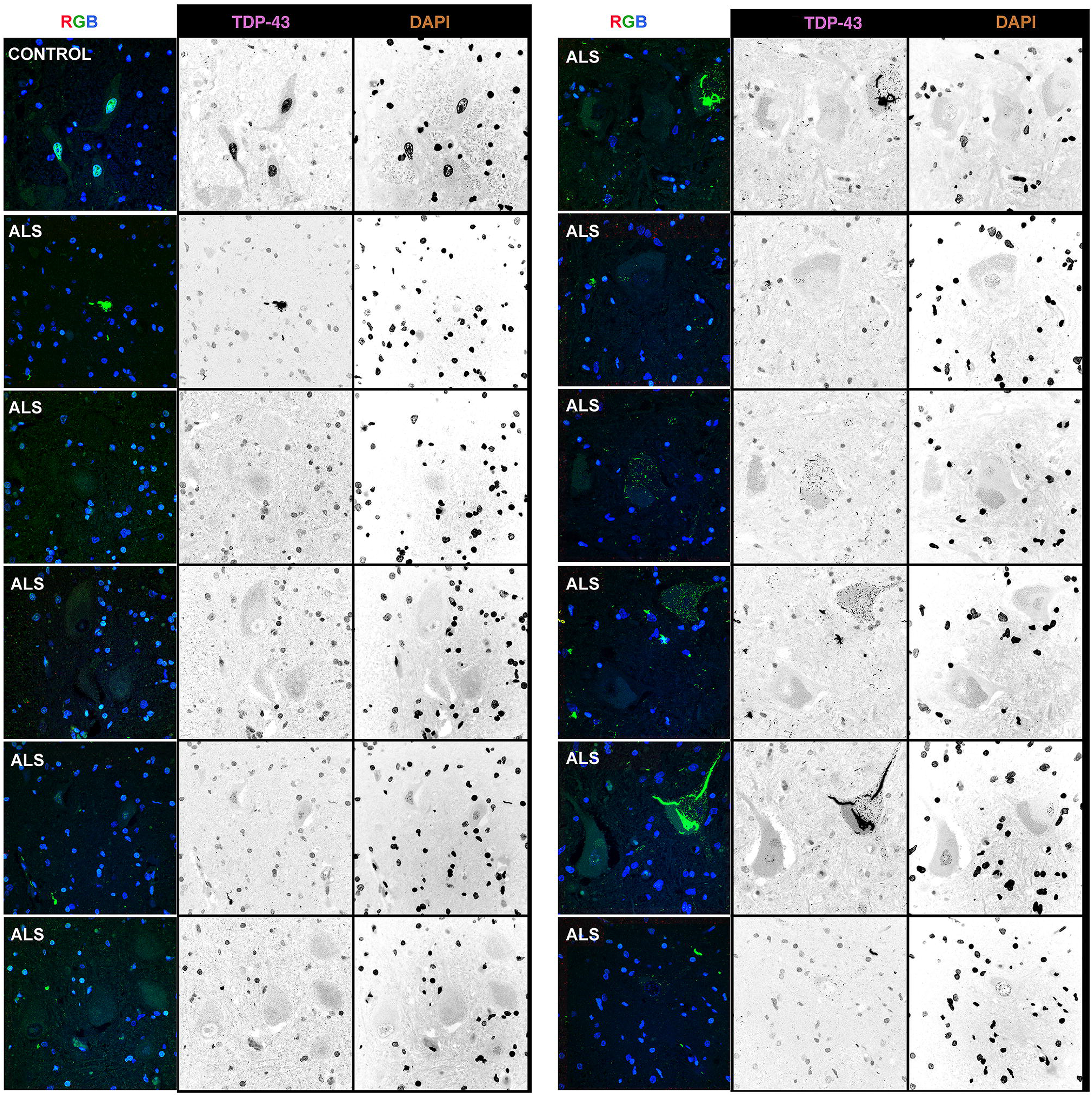
Different examples of TDP-43 pathology in neurons and glial cell in controls and ALS motor neurons.

**Supplementary Figure 9.**
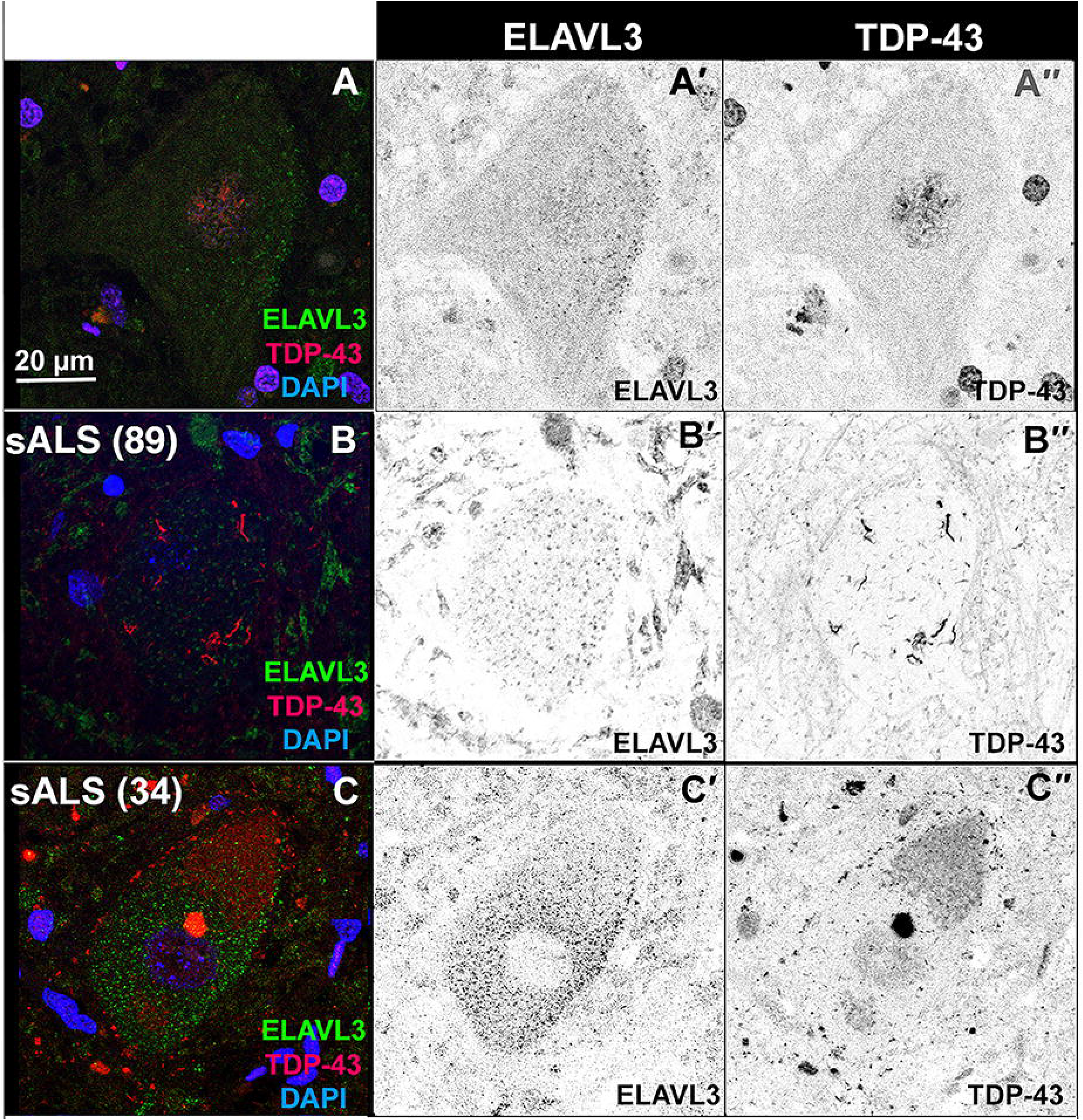
ELAVL3 inclusions do not co-localized with TDP-43 aggregates. A. Control motor neuron has nuclear ELAVL3 and nuclear TDP-43. B-C. Cytoplasmic inclusions of ELAVL3 and TDP-43 in motor neurons do not co-localize. [Scale bar 10μm.]

**Supplementary Figure 10.**
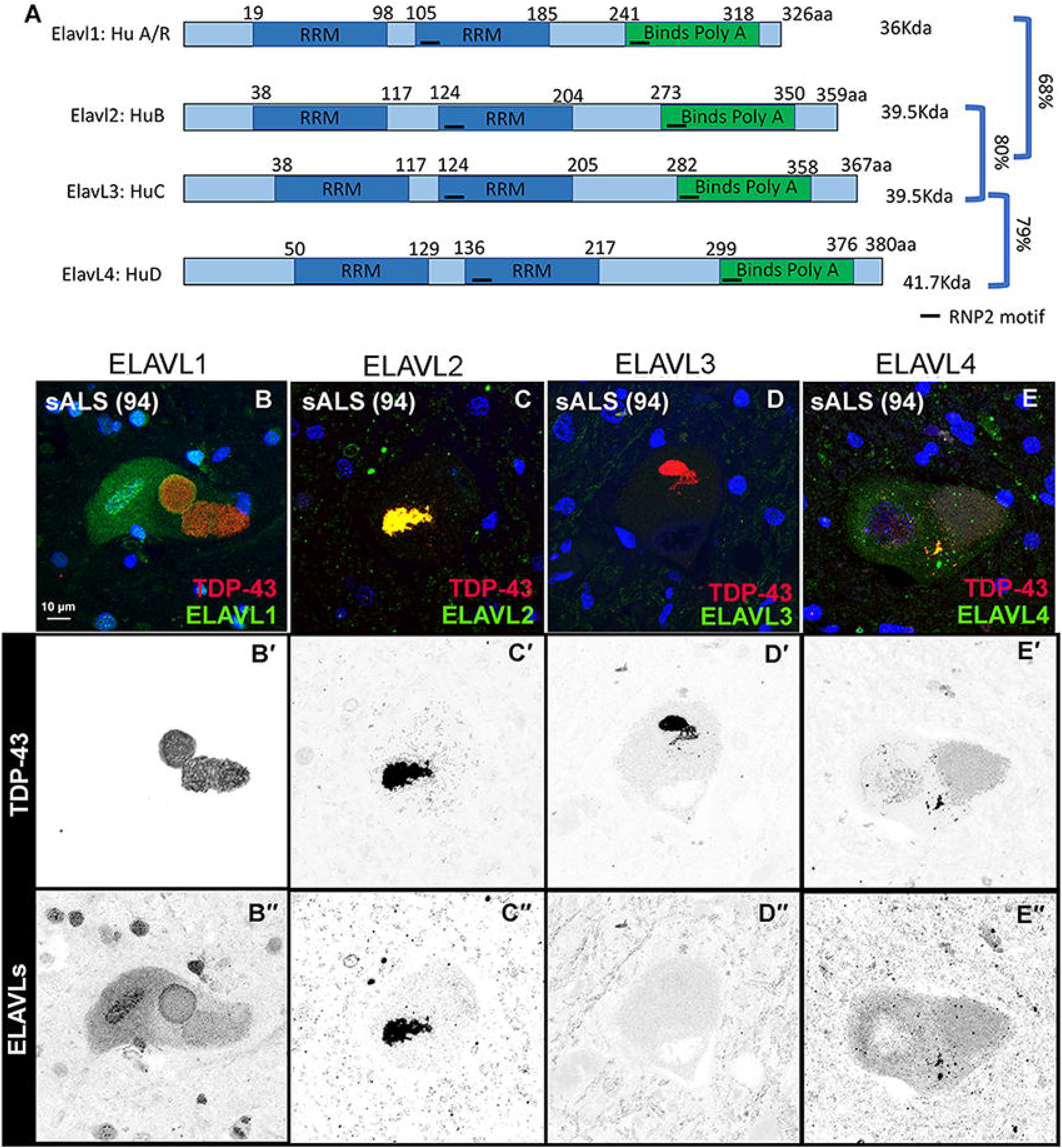
ELAVLs protein behave different within the same nervous system. A. Scheme of DNA sequence homology between different ELAVL3 family members. B. Staining of ELAVL1, 2, 3 and 4 in the same patient at contiguous sections. ELAVL1 is seen encapsulating TDP-43 aggregate. ELAVL2 co-localized with TDP-43 aggregates. ELAVL3 is nuclear depleted. ELAVL4 is depleted in the nucleus and aggregated in the cytoplasm where it partially co-localizes with TDP-43.

**Supplementary Figure 11.**
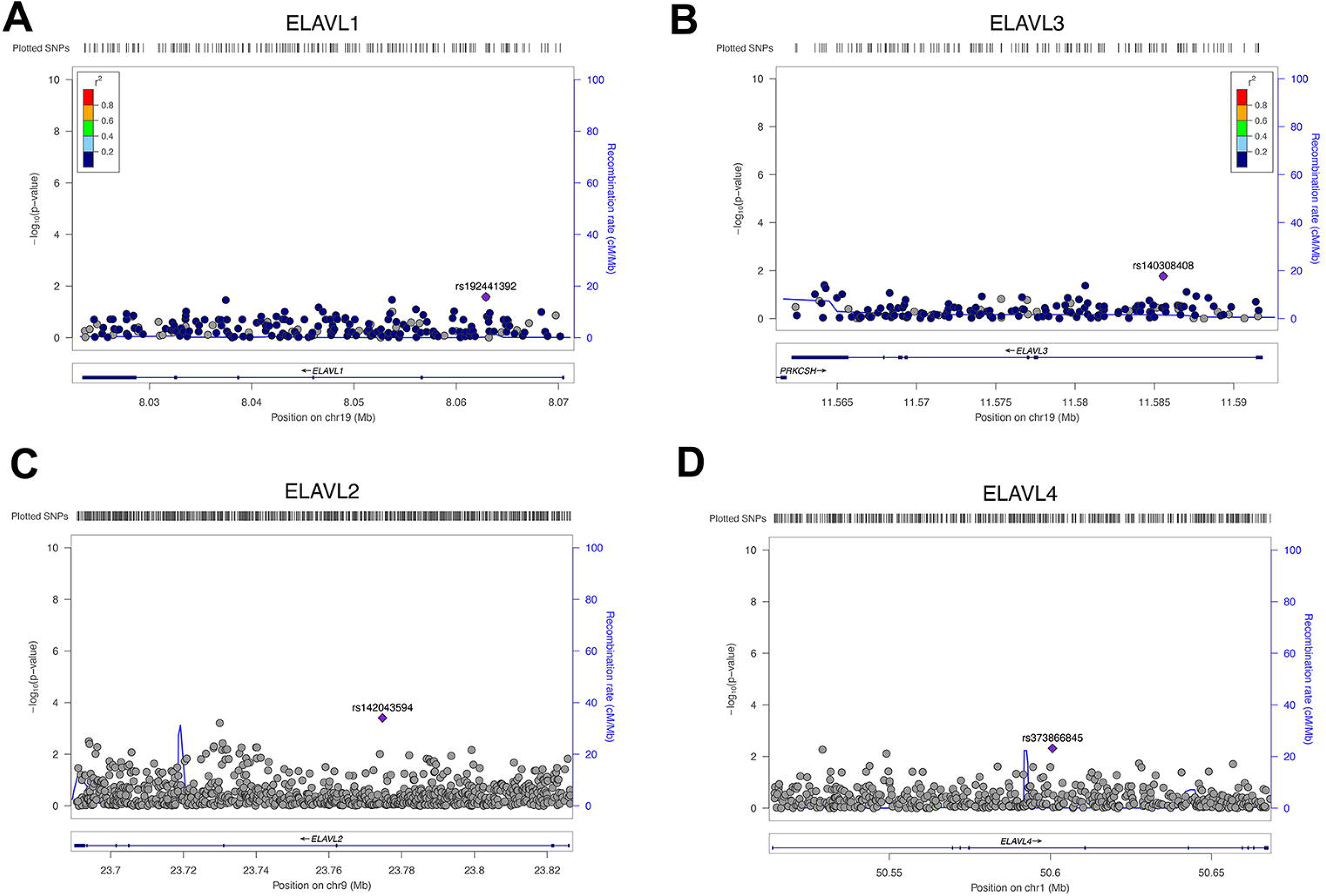
Genetic variants of ELAVL family members were not found to be associated with high risk ALS.

## Acknowledgements

This research was supported by grants from National Institute of Neurological Disorders and Stroke (NIH R01NS088578 and R21NS121805), UCSD Microscopy Core (NINDS NS047101), ALS Association, Target ALS, Microsoft Research, Pam Golden and the Kraatz Family/Nicholas Martin Jr. Family Foundation. We thank Target ALS CNS biorepository, the patients, and their families for their generous contribution to this research. We also would like to thank the Nikon Imaging Center at UC San Diego for assistance with microscopy and analysis and UCSD School of Medicine Microscopy Core (Grant P30 NS047101).

